# Presynaptic Ca_V_2.1 calcium dependent facilitation is essential for faithful auditory information transfer

**DOI:** 10.64898/2026.04.24.720705

**Authors:** Mohammed Al-Yaari, Jianing Li, Christian Keine, Bryce Hunger, Marlan R. Hansen, Samuel M. Young

## Abstract

Activity-dependent modulation of presynaptic voltage-gated Ca^2+^ channels (Ca_V_2) regulates Ca^2+^ influx to control neuronal circuit output. Although Ca_V_2.1 can undergo robust Ca^2+^ dependent facilitation (CDF), whether it occurs in native central nervous system neuronal circuits is disputed. Accurate auditory information processing requires precise and reliable synaptic transmission at high activity rates in the auditory brainstem. To determine if Ca_V_2.1 CDF is a key regulator of high-fidelity synaptic transmission, we expressed Ca_V_2.1 splice variants capable (Ca_V_2.1 37a) or incapable (Ca_V_2.1 37b) of CDF at the calyx of Held presynaptic terminal in the auditory brainstem. We found no difference in basal Ca_V_2.1 currents or synaptic transmission. However, Ca_V_2.1 37b terminals lacked CDF, synaptic facilitation and had a decreased reliability and precision of postsynaptic action-potential firing. Additionally, the Wave III amplitude of the auditory brainstem responses was reduced. We propose that Ca_V_2.1 CDF is essential for accurate auditory information processing.

## Introduction

Accurate sound encoding requires synapses in the lower auditory brainstem to drive and sustain temporally precise action potential (AP) firing over rapid and large fluctuations in AP firing rates of upwards to one kHz^1–4^. However, the number of synaptic vesicles (SVs) available for AP-evoked release, the readily releasable pool (RRP), is limited^5^. Therefore, extremely rapid SV release and RRP replenishment is needed for accurate auditory information encoding^4,6–8^. Activity-dependent increases in presynaptic [Ca^2+^] increase synaptic strength and SV replenishment speeds which modulates the reliability and precision of synaptic transmission^8–12^. Therefore, the molecular mechanisms regulating activity-dependent changes in presynaptic [Ca^2+^] are essential to understanding how sound information is accurately encoded.

The magnitude of Ca^2+^ influx through presynaptic voltage-gated calcium channels (Ca_V_) can change in an activity-dependent manner to enhance or decrease synaptic strength^11–13^. In response to activity, Ca^2+^ currents can undergo calcium-dependent facilitation (CDF) and calcium-dependent inactivation (CDI)^14^. In central synapses, Ca_V_2.1 channels are the dominant subtype mediating AP-evoked release^15^. Among the Ca_V_2 family, robust CDF is unique to Ca_V_2.1 and has been proposed to be an important component of synaptic facilitation^16^. A multitude of structure/function studies have demonstrated that the EF-hand, IQ-like domains, and Calmodulin binding Domain (CBD) in the Ca_V_2.1 α_1_ subunit are essential determinants that act in concert to promote Ca_V_2.1 CDF^17,18^.

In the lower auditory brainstem, the presynaptic terminals in the initial synaptic connections encoding sound localization are Ca_V_2.1 exclusive and undergo synaptic facilitation in an activity-dependent manner^15^. The calyx of Held/medial nucleus of the trapezoid body (MNTB) synapse is a key synapse located in the superior olivary complex (SOC) that encodes sound localization^19–22^. The calyx of Held is an axosomatic presynaptic Ca_V_2.1 exclusive terminal that is the sole input driving AP firing in the MNTB with high temporal precision and nearly fail safe synaptic transmission^1,10^. At the calyx of Held, Ca_V_2.1 CDF leads to activity-dependent increases in presynaptic [Ca^2+^] as loss of Ca_V_2.1 results in an absence of activity-dependent increases in presynaptic [Ca^2+^] levels^23,24^. Furthermore, direct presynaptic recordings at the immature calyx (prehearing) at low external [Ca^2+^] and room temperature (25°C), revealed activity dependent on Ca^2+^ increases^25^. Therefore, it is hypothesized that Ca_V_2.1 CDF is a key regulator of accurate sound information encoding^1,4,26,27^.

Due to disparate results, the contribution of Ca_V_2.1 CDF to synaptic facilitation and the efficacy of synaptic transmission is highly disputed^28–31^. The Ca_V_2.1 α_1_ subunit exon 37 codes for the EF-hand an essential element for Ca_V_2.1 CDF^32,33^. Due to mutually exclusive splicing, the Ca_V_2.1 α_1_ subunit contains Exon 37a (37a) or Exon37b (37b) with Ca_V_2.1 37a capable of CDF, and Ca_V_2.1 37b incapable^32,34^. In dissociated Purkinje cells at room temperature and 5 mM [Ca^2+^]_ext_, Ca_V_2.1 37a led to significant activity dependent increases in [Ca^2+^] while Ca_V_2.1 37b did not^34^. However, Ca_V_2.1 37b overexpression in cultured primary hippocampal neurons led to significant increases in synaptic facilitation^31^. The Ca_V_2.1 α_1_ subunit EF-hand IQ channel mutant global knock in mouse (Ca_V_2.1 IM-AA) which ablates Ca_V_2.1 CDF resulted in significant reductions in synaptic facilitation in native Schaffer Collateral (SC)-CA1 synapses, and CA3/PV interneuron synapses and at the mouse neuromuscular junction (NMJ)^29,30,35^. In contrast, at physiological temperatures and [Ca^2+^]_ext_ synaptic facilitation was not reduced in Ca_V_2.1 IM-AA hippocampal CA3/CA1 synapses or PF/PC and PC/DCN cerebellar synapses^28^. Furthermore, there was little activity dependent Ca_V_2.1 CDF up to 100 Hz in the soma of dissociated Purkinje cells^28^. Therefore, it was concluded that under physiological conditions in these synapses Ca_V_2.1 CDF makes little contribution to synaptic facilitation^28^.

To directly determine if Ca_V_2.1 CDF is required for reliable and temporally precise synaptic transmission for encoding auditory information, we used viral vectors to express either the Ca_V_2.1 37a or Ca_V_2.1 37b α_1_ subunit while ablating endogenous Ca_V_2.1 α_1_ subunit at the calyx of Held in a Ca_V_2.1 conditional knockout (CKO) mouse model^36^. Direct presynaptic recordings at physiological temperatures and [Ca^2+^]_ext_ revealed no difference in AP-evoked release synaptic transmission between Ca_V_2.1 37a or Ca_V_2.1 37b calyces. Although there was no difference in initial Ca_V_2.1 currents, only Ca_V_2.1 37a calyces had activity-dependent increases in [Ca^2+^]. Furthermore, at Ca_V_2.1 37b calyx/MNTB synapses we found an absence of synaptic facilitation, increased synaptic depression, higher AP failure rates, and reduced AP temporal precision in an activity-dependent manner. Auditory brainstem responses (ABR) from animals with calyces expressing Ca_V_2.1 37b exhibited significant reductions in wave III amplitudes corresponding to reduced activity in the SOC. Taken together, we propose that presynaptic Ca_V_2.1 CDF is essential for accurate auditory information processing.

## Results

### Ca_V_2.1 Exon37 does not regulate presynaptic Ca^2+^ levels and the basal synaptic transmission

To probe the importance of Ca_V_2.1 CDF for temporally precise and reliable synaptic transmission in the auditory brainstem, we created Helper-dependent Adenoviral vectors (HdAd) that expressed native full length mouse Ca_V_2.1 α_1_ subunit splice variants with either exon 37a (Ca_V_2.1 37a) or exon 37b (Ca_V_2.1 37b) under the 470 bp human synapsin promoter transgene expression cassette to drive Ca_V_2.1 expression^36^ **(Fig 1A**). Subsequently, using established techniques, we co-injected HdAd Cre into the P1 cochlear nucleus (CN) of a *Cacna1A* CKO mouse and either HdAd Ca_V_2.1 37a or HdAd Ca_V_2.1 37b to create a Ca_V_2.1 37a or Ca_V_2.1 37b exclusive calyx of Held^37^ (**Fig 1 A, B**). Subsequently, all experiments were performed at the calyx of Held/MNTB synapse from P16 and onwards which is the developmental time point where the calyx is Ca_V_2.1 exclusive, mature and the neuronal circuit properties are considered “adult-_like”10,38-40._

**Figure 1.**
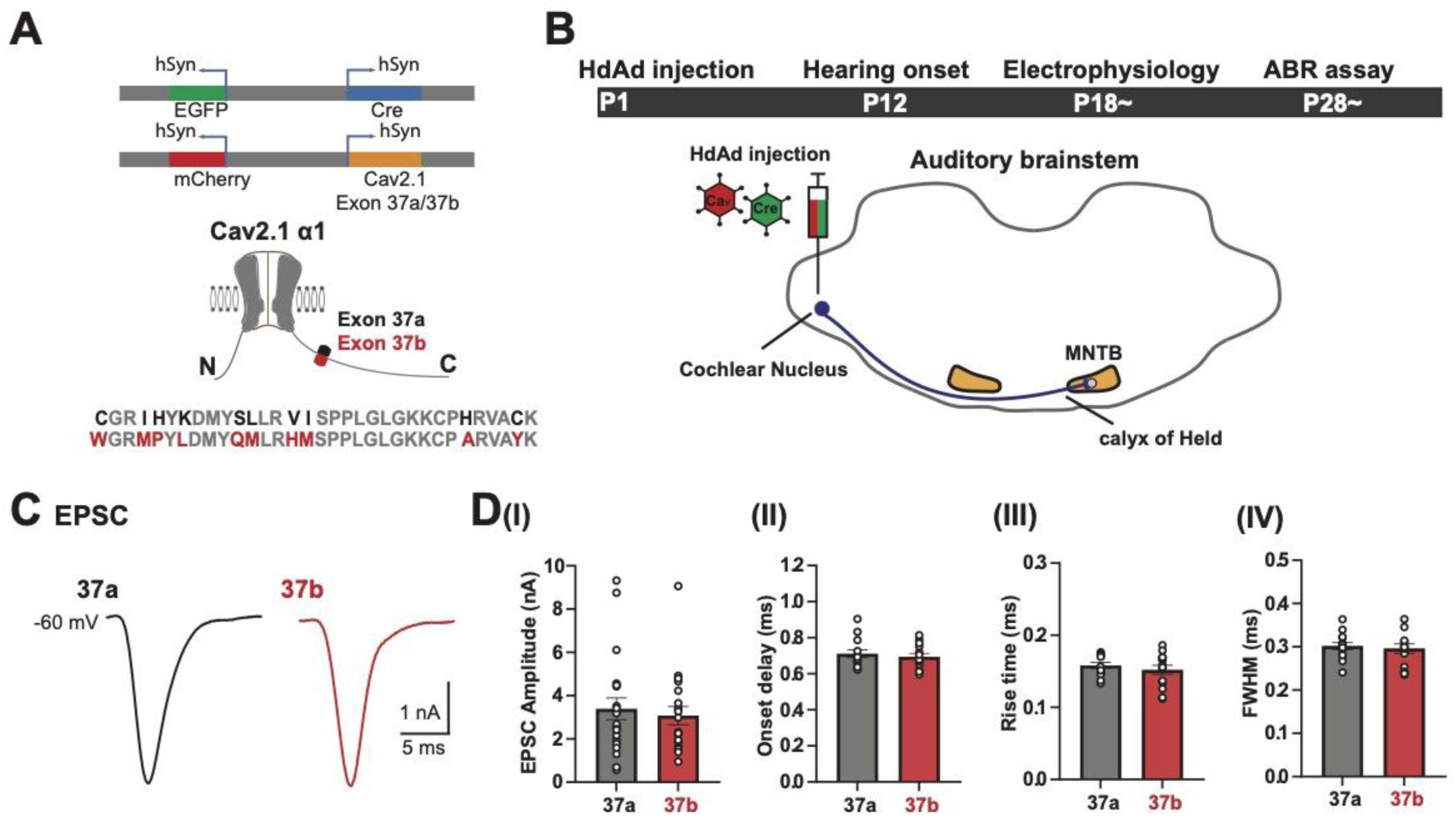
Ca_V_2.1 Exon37 a/b isoforms do not regulate the basal synaptic transmission. A: A cartoon showing Ca_V_2.1 α_1_ subunit with two different splice variants in the C-terminus EF-hand (Ca_V_2.1 37a in black and Ca_V_2.1 37b in red) (bottom). Schematic of the viral constructs used, expressing either Cre + EGFP and Ca_V_2.1 37a or 37b + mCherry, respectively, driven by the synapsin promoter (top left). **B:** The experimental timeline (top), a schematic auditory brainstem showing the stereotaxic surgery at P1 with injecting HdAd vectors expressing Cre +EGFP and Ca_V_2.1 37a or 37b mCherry into the cochlear nucleus, and the fibers arise from Globular Bushy cells (GBC) to the MNTB neurons (blue color) (bottom right). **C**: Representative traces of postsynaptic whole-cell recording from principal MNTB neurons voltage-clamped at −60 mV and afferent fibers (calyces) were stimulated by a bipolar electrode. **D:** Comparison of the characteristics of synaptic strength and kinetics, (EPSC amplitude (I), onset delay (II), rise time (III) at 10-90%, and FWHM at 50% (IV) collected from MNTB neurons; *37a* n = 22, 8 animals; *37b* = 22, 11 animals. EPSC amplitude; 37a: 3.14 ± 0.50, 37b: 3.16 ± 0.4 nA, onset delay 37a: 0.72 ± 0.02, 37b: 0.7 ± 0.16 ms, rise time; 37a: 0.16 ± 0.004, 37b: 0.15 ± 0.01 ms, FWHM; 37a 0.31±0.01, 37b: 0.29 0.01 ms, mean ± SEM.

The Ca_V_2.1 exon 37a has been proposed to regulate presynaptic Ca_V_2.1 abundance and coupling, the distance between the Ca_V_2.1 channel and SV calcium sensor^31,33^. The mature calyx of Held is a nanodomain release synapse with SVs coupled to Ca_V_2.1 channels at ∼25nm and AP evoked release is highly sensitive to changes in coupling distance and Ca^2+^ current levels^41–43^. To indirectly test if there were differences in Ca^2+^ current levels or coupling distances, we carried out afferent fiber stimulation at 0.1 Hz and measured AMPAR currents at MNTB neurons innervated by either Ca_V_2.1 37a and Ca_V_2.1 37b calyces. Our analysis revealed no difference in the AP-evoked EPSC amplitude, EPSC rise time, FWHM or onset delay (**Fig 1D**). Taken together, our data show that the Ca_V_2.1 EF-hand does not regulate presynaptic Ca_V_2.1 abundance or coupling.

### Presynaptic Ca^2+^ currents in Ca_V_2.1 37a calyces but not Ca_V_2.1 37b calyces undergo AP-evoked activity dependent increases at physiological temperatures and [Ca^2+^]_ext_

The magnitude of Ca_V_2.1 CDF is dependent on the internal [Ca^2+^]^14^. At the soma in dissociated Purkinje cells and hippocampal neurons at physiological temperatures and [Ca^2+^]_ext_ (∼1.5 mM) activity-dependent Ca^2+^ increases were considered negligible^28^. In contrast, direct presynaptic recordings at the immature calyx (P9-P11, prehearing) revealed activity-dependent Ca^2+^ increases at 1.0 mM [Ca^2+^]_ext_ at room temperature (25°C)^25^. Since mature calyx AP-evoked Ca^2+^ currents at physiological temperatures are smaller than at the immature calyx^44^, it was possible this do not occur at mature calyces. To test this, we performed whole cell patch clamp recordings at Ca_V_2.1 37a or Ca_V_2.1 37b calyces at physiological temperature and [Ca^2+^]_ext_ (∼35°C and 1.2 mM) (**Fig 2**). Using a mature AP-evoked waveform (**Fig 2A**), we measured activity-dependent changes in presynaptic Ca^2+^ current in response to 50 AP at three stimulation frequences (100 Hz, 300 Hz, and 500 Hz) that are typical firing rates for encoding auditory information. Our analysis revealed no difference in initial Ca_V_2.1 current levels (**Fig 2B**) indicating no difference in Cav2.1 expression levels. However, we found activity-dependent increases in presynaptic Ca^2+^ currents in the Ca_V_2.1 37a calyces at 300 and 500Hz but a complete absence of activity-dependent Ca^2+^ current increases in Ca_V_2.1 37b calyces (**Fig 2C, 2D**). In addition, we performed presynaptic recordings of wild-type calyces and found robust CDF (**Fig S1**). Based on these data, we conclude that Ca_V_2.1 CDF occurs at the mature calyx and results in activity-dependent increases in presynaptic Ca^2+^ current levels under physiological conditions.

**Figure 2.**
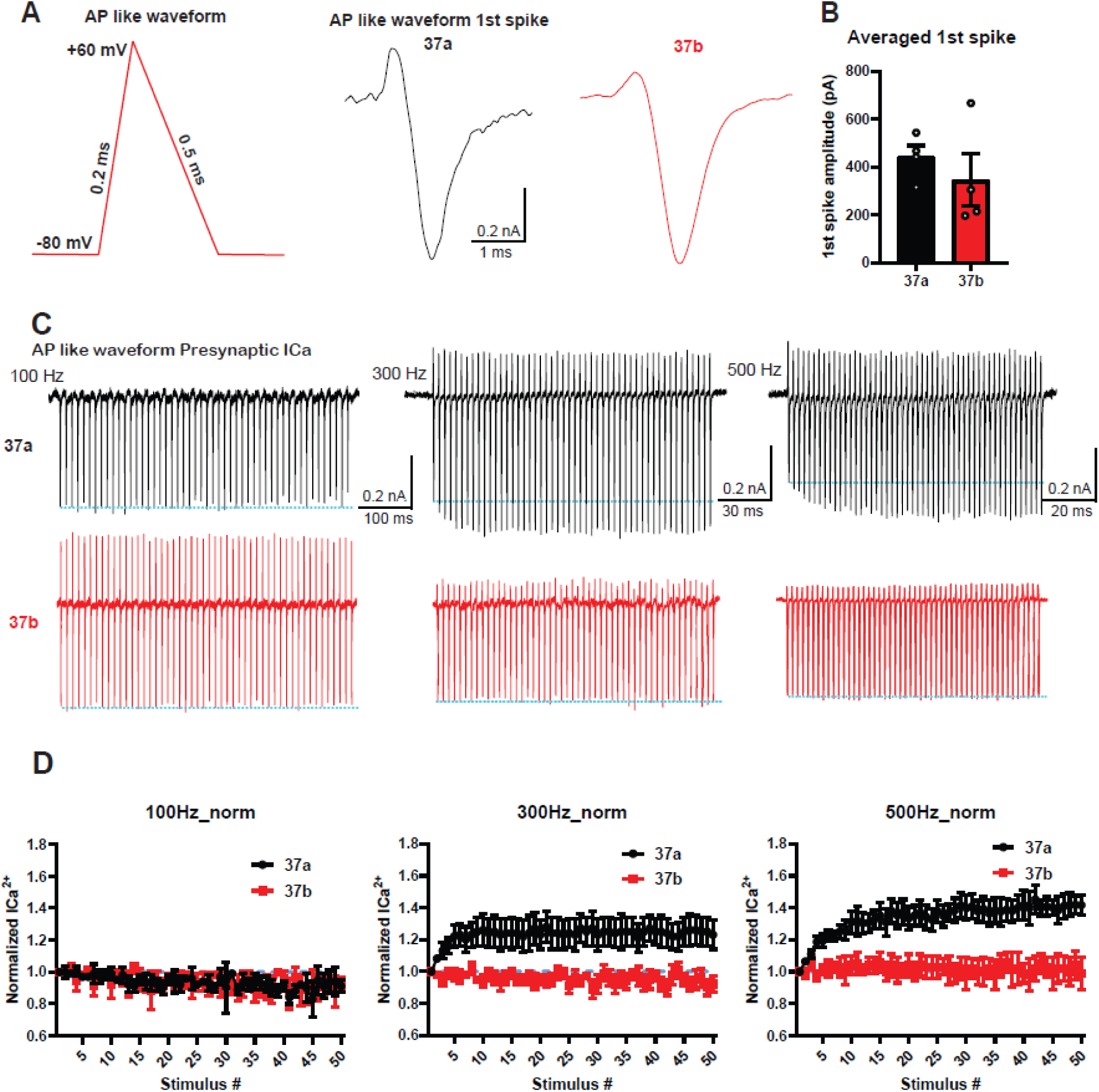
Ca_V_2.1 37a but not Ca_V_2.1 37b undergoes CDF in a frequency dependent manner at physiological temperature and external [Ca^2+^] A: Representative traces of presynaptic AP like waveform (right) at calyces expressing either Ca_V_2.1 37a (black) or Ca_V_2.1 37b (red) with a depolarization ramp at 60 mV; the ramp depolarization has rising time (0.2ms) and falling phase (0.5ms) (left). **B:** Bar graph showing averaged amplitude of 1^st^ spike in the AP like stimulation. Each data point is the average of 1^st^ spike of 100Hz, 300Hz and 500Hz AP like stimulation per cell. 37a, 441.8±47.48 pA, n=4, 4 animals. 37b, 345.5±109.6, n=4, 4 animals, mean ± SEM, p=0.4509, unpaired t-test; **p < 0.01, ***p < 0.001, ****p < 0.0001. **C:** Presynaptic recordings of AP like waveform train (100, 300, 500 Hz) at calyces expressing Ca_V_2.1 37a (black) or 37b (red). **D:** Graphs of Ca^2+^ dependent facilitation, Ca^2+^ current amplitudes are normalized to the 1^st^ spike and plotted against stimulus number at different frequencies (100, 300, 500 Hz). 100 Hz: 37a n=4, 4 animals. 37b n=2, 2 animals, p=0.3885. 300 Hz: 37a n=4, 4 animals. 37b n=4, 4 animals, p<0.0001. 500 Hz: 37a n=4, 4 animals. 37b n=4, 4 animals, p<0.0001, paired t-test; **p < 0.01, ***p < 0.001, ****p < 0.0001.

### Absence of Ca_V_2.1 CDF abolished synaptic facilitation and decreased the steady-state levels under *in vivo*-like conditions

Activity-dependent increases in AP firing lead to multiple forms of short-term plasticity which include short-term synaptic facilitation at the calyx of Held/MNTB synapse both *in vitro* and *in vivo* conditions^8^. However, *in vivo* background spontaneous activity tonically depresses the mature calyx of Held, which reduces the magnitude of short-term facilitation and depression^7,10^. Therefore, to test if Ca_V_2.1 CDF was a key regulator of short-term plasticity we performed afferent fiber stimulation of transduced calyces expressing Ca_V_2.1 37a or Ca_V_2.1 37b and recorded

AMPAR currents at the MNTB at physiological temperature and [Ca^2+^]. To mimic *in vivo* like conditions, we used a 2 second conditioning train, 60 APs 30 Hz (conditioning), followed by test trains of 50 APs at different stimulus frequencies, 100, 300, and 500 Hz (test). The 2s conditioning train stimulus frequency was chosen as this represents the average spontaneous firing rate *in vivo*^7^, while the stimulation frequencies in test trains are typical firing rates during the encoding of auditory information.

Analysis of the conditioning train from Ca_V_2.1 37a and Ca_V_2.1 37b calyces revealed similar initial EPSC amplitudes, paired pulse ratio (PPR), short-term plasticity, and steady state levels (**Fig S2**). However, test train stimuli at the different stimulation frequencies revealed differences in frequency-dependent plasticity and steady-state EPSC amplitudes (**Fig 3**). Although test trains from Ca_V_2.1 37a and 37b calyces had similar initial EPSC amplitudes at the start of the test train, normalization to the initial EPSC1 amplitude revealed Ca_V_2.1 37a synapses underwent frequency-dependent facilitation early in the train followed by synaptic depression, while Ca_V_2.1 37b synapses underwent frequency-dependent depression through the entire train. Additionally, PPR analysis confirmed that Ca_V_2.1 37a underwent paired pulse facilitation (PPF) while Ca_V_2.1 37b calyces did not at any_frequencies tested, ((**Fig 3D** ; PPR; (37a 100 Hz: 1.167 ± 0.08, 37b: 0.85 ± 0.06), (37a 300 Hz: 1.33 ± 0.081, 37b: 1.046 ± 0.076), (37a 500 Hz 1.44 ± 0.134, 37b: 0.98 ± 0.054)). Finally, there were differences between the normalized steady state EPSCs at all frequencies tested. EPSCss: (100 Hz 37a; 1.03 ± 0.09 n=10, 37b; 0.71 ± 0.05 n=11 p= 0.01, 300 Hz 37a; 0.93 ± 0.1 n=10, 37b; 0.64 ± 0.1 n=12 p= 0.05, 500 Hz 37a; 0.9 ± 0.2 n=8, 37b; 0.37 ± 0.05 n=11 p=0.0.1. However, back extrapolation of the cumulative EPSCs did not reveal any differences in replenishment rates although there appeared an activity dependent increase in the cumulative EPSC (**Fig S3A,C**).

**Figure 3.**
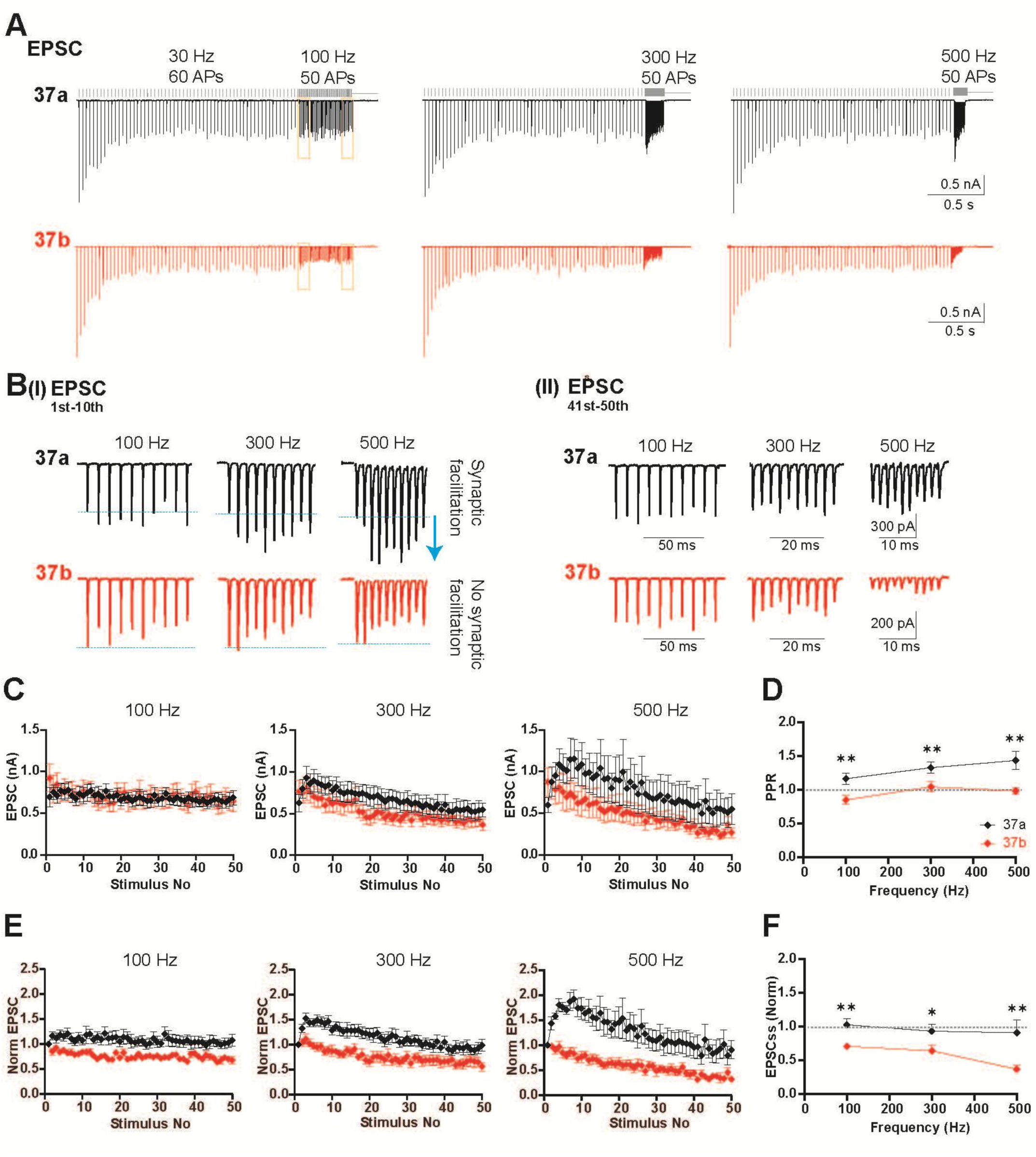
Lack of Ca^2+^ CDF abolished synaptic facilitation and decreased the steady-state level of EPSC *in vivo* like responses. **A**: EPSCs traces of test trains (50 Action Potentials (APs) at different stimulus frequencies (100, 300, 500 Hz) preceded of conditioned trains (60 APs) at 30 Hz for ∼ 2 s. EPSCs are recorded from MNTB neurons receiving calyces expressing either Ca_V_2.1 37a (top, black traces) or Ca_V_2.1 37b (bottom, red traces). **B:** Zoomed in traces of EPSC₁–EPSC₁₀ or EPSC₄₁–₅₀ showed on the bottom of the representative traces in B(I) and B(II) respectively. 37b neurons showed no synaptic facilitation in the early stimuli (∼20 ms) of high frequency stimulation (300, 500 Hz) and a lower steady-state level in comparison to those of 37a neurons. **C**: Graphs of absolute EPSC values against stimulus numbers at different stimulus frequencies (100, 300, 500 Hz). Synaptic facilitation is abolished in high frequencies (300, 500 Hz) of 37b neurons, and the level of steady state is lower than that of 37a neurons. **D:** Paired-pulse ratio (PPR) against stimulus frequencies (100, 300, 500 Hz). MNTB neurons receiving 37a calyces showed paired-pulse facilitation while those with 37b calyces showed no facilitation. Statistical analysis showed a significant difference between Ca_V_2.1 exon37a and 37b among all tested stimulus frequencies. **E:** Graphs of normalized EPSCs to the 1^st^ EPSC amplitude against stimulus numbers at different stimulus frequencies (100, 300, 500 Hz). The synaptic facilitation is more prominent in all tested frequencies, particularly (300, 500 Hz) of 37a neurons. **F:** Average EPSC 41^st^- 50^th^ (EPSCss) normalized to the 1^st^ EPSC against stimulus frequencies. The level of steady state almost alike to the 1^st^ EPSC of Ca_V_2.1 37a synapses, however, it usually gets lower than the 1^st^ EPSC in all tested frequencies of Ca_V_2.1 37b synapses and showed significant statistical difference between the two groups in all frequencies. PPR: PPR: 100 Hz 37a; 1.2 ± 0.08 n=10, 37b; 0.85 ± 0.06 n=11 p=0.01, 300 Hz 37a; 1.3 ± 0.08 n=10, 37b; 1 ± 0.07 n=12 p= 0.01, 500 Hz 37a; 1.4 ± 0.13 n=8, 37b; 0.98 ± 0.05 n=11 p= 0.01. EPSCss: 100 Hz 37a; 1.03 ± 0.09 n=10, 37b; 0.71 ± 0.05 n=11 p= 0.01, 300 Hz 37a; 0.93 ± 0.1 =10, 37b; 0.64 ± 0.1 n=12 p= 0.05, 500 Hz 37a; 0.9 ± 0.2 n=8, 37b; 0.37 ± 0.05 n=11 p=0.01. unpaired t-test; **p < 0.01, ***p < 0.001, ****p < 0.0001

To determine how synaptic plasticity in the tonically depressed calyx compared to the “rested” calyx, we performed similar train stimuli without conditioning (**Fig 4**). Similar to the tonically depressed calyces, Ca_V_2.1 37a calyces exhibited frequency-dependent facilitation and PPF but not Ca_V_2.1 37b calyces. Comparable to tonically depressed calyces, there were not differences in the replenishment rates, however unlike the tonically depressed calyces there was no activity dependent increase the cumulative EPSCs (**Fig S3C,D**). Finally, to rule out compensation by Ca_V_2.2 channels, we expressed Ca_V_2.1 37a or Ca_V_2.1 37b on the Ca_V_2.1/ Ca_V_2.2 null calyx background. Results of our analyses revealed similar phenotypes (**Fig S4**). Based on these results, we conclude that Ca_V_2.1 CDF is essential for frequency-dependent facilitation and regulates steady-state levels at the calyx of Held/MNTB synapse.

**Figure 4.**
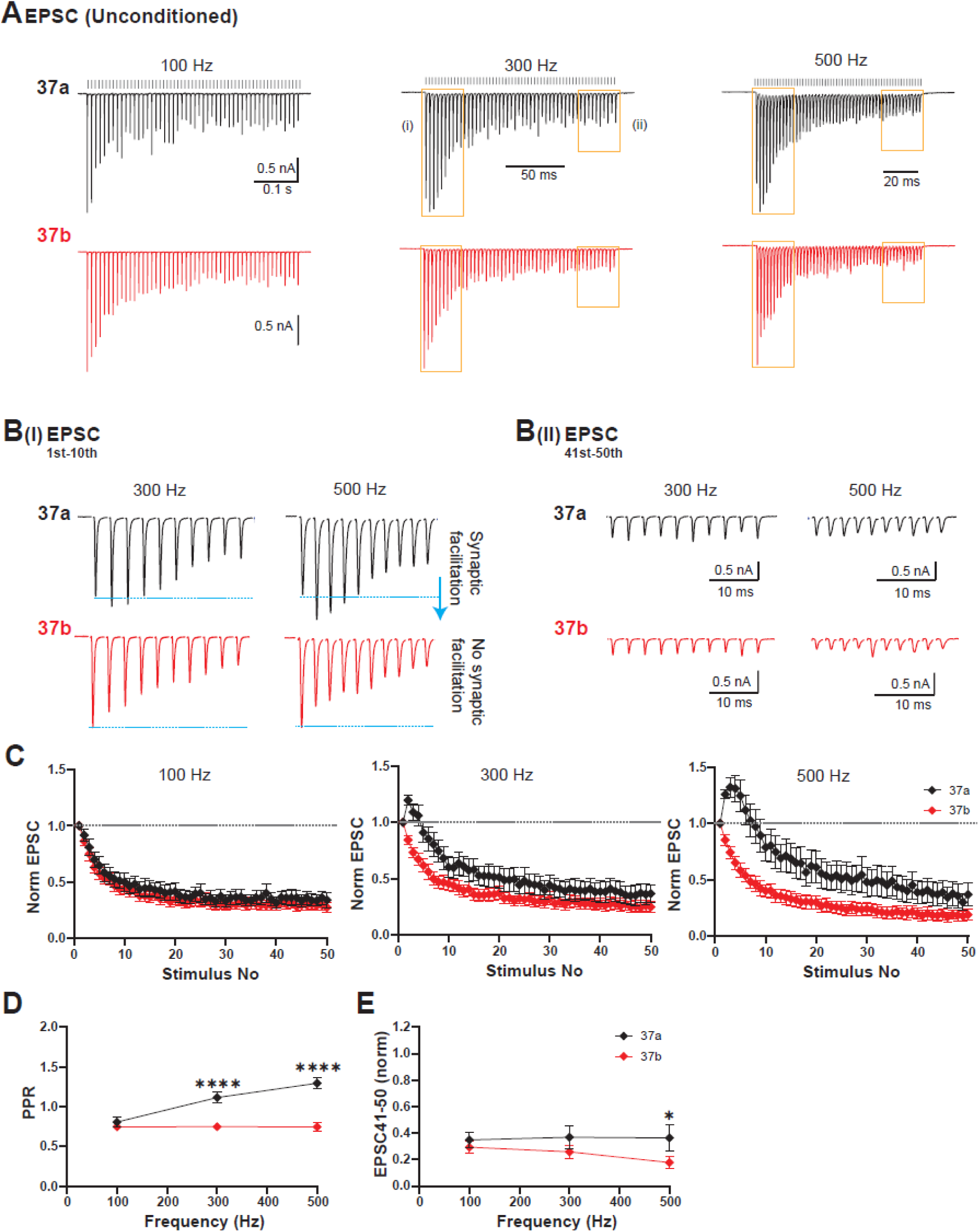
Lack of Ca^2+^ current facilitation abolished synaptic facilitation and decreased the steady-state level of EPSC in conventional electrical stimulation. A: Representative EPSC traces evoked by test trains of 50 APs at 100, 300, and 500 Hz without preceding conditioning trains. **B:** Expanded views of EPSC₁–EPSC₁₀ (**B-I**) and EPSC₄₁–EPSC₅₀ (**B-II**) taken from the traces shown in panel **A**. Neurons receiving 37b calyces exhibited no synaptic facilitation during the early phase of high-frequency stimulation (300 and 500 Hz) and reached a lower steady-state level compared with neurons receiving 37a calyces, which is consistent with our observations in Figure 3 using *in vivo*–like conditioning (preceded by 60 APs at 30 Hz). **C:** Plots of normalized EPSC amplitude versus stimulus numbers at 100, 300, and 500 Hz. Synaptic facilitation was abolished at high frequencies (300 and 500 Hz) in neurons receiving 37b calyces, and their steady-state levels were lower than those of 37a neurons. **D:** PPR as a function of stimulation frequency. MNTB neurons receiving 37a calyces showed paired-pulse facilitation, particularly at high frequencies (300 and 500 Hz), whereas neurons receiving 37b calyces showed little to no facilitation. **E:** Average steady-state EPSCs (EPSC₄₁–₅₀) normalized to the 1^st^ EPSC plotted against stimulating frequency. Steady-state amplitude decreased in a frequency-dependent manner to <50% of the 1^st^ EPSC and was significantly lower in neurons receiving 37b calyces, particularly at 500 Hz. PPR: 100 Hz 37a; 0.9 ± 0.04 n=14, 37b; 0.85 ± 0.04 n=14 p=0.3, 300 Hz 37a; 1.2 ± 0.03 n=17, 37b; 0.84± 0.04 n=16, p < 0.0001, 500 Hz 37a; 1.25 ± 0.03 n=15, 37b; 0.85± 0.04 n=18, p<0.0001. EPSCss: 100 Hz 37a; 0.35 ± 0.06 n=13, 37b; 0.3 ± 0.04 n=13 p= 0.5, 300 Hz 37a; 0.37 ± 0.09 n=14, 37b 0.26± 0.06 n=16 p= 0.2, 500 Hz 37a; 0.3 ± 0.07 n=12, 37b; 0.18 ± 0.05 n=16 p=0.04. unpaired t-test; **p < 0.01, ***p < 0.001, ****p < 0.0001.

### Ca_V_2.1 CDF is critical for synaptic reliability and temporal fidelity of AP spiking in the MNTB

The calyx of Held/MNTB synapse is considered fail-safe relay due to its ability to maintain AP-spiking up to 1 kHz^1^. Therefore, we examined how the absence of CDF impacted AP-firing reliability in the MNTB neuron (**Fig 5**). To do so, we performed loose cell-attached recordings from MNTB neurons using the stimulation paradigm in **Fig 3**, (2s 30Hz conditioning train, followed by test train). Kynurenic acid and AP-5 were not included in the external solution. Firing probability was recorded as presence or absence of postsynaptic AP in a single neuron and the averaged across all neurons (See Materials and Methods). Subsequently, we plotted the firing probability against stimulation number. We found that there were no spike failures in conditioning trains for either Ca_V_2.1 37a and Ca_V_2.1 37b synapses (**Fig S5).** However, while both Ca_V_2.1 37a and Ca_V_2.1 37b exhibited increases in postsynaptic spiking failures in a frequency-dependent manner, the effect was more severe in Ca_V_2.1 37b synapses. In particular, Ca_V_2.1 37b synapses showed occasional failures at 100 Hz, while 37a calyces did not. This difference became more pronounced at 300 Hz and 500 Hz stimulation frequency, with Ca_V_2.1 37b showing significant impairment of firing probability even early in the train (**Fig. 5C**). To correlate the firing probability during steady state EPSC we averaged the firing probability of the steady-state (Spikes 41-50) and compared the steady state firing probability between Ca_V_2.1 37a and Ca_V_2.1 37b. We found Ca_V_2.1 splice variant also impacted particularly at 500 Hz: spikes 41-50 100 Hz 37a; 1 ± 0.002, 37b; 0.96 ± 0.03, p=0.009, 300 Hz 37a; 0.90 ± 0.05, 37b; 0.78 ± 0.05, p=0.03, 500 Hz 37a; 0.62±0.046, 37b; 0.44±0.05, p=0.04). (**Fig 5B,C)**

**Figure 5.**
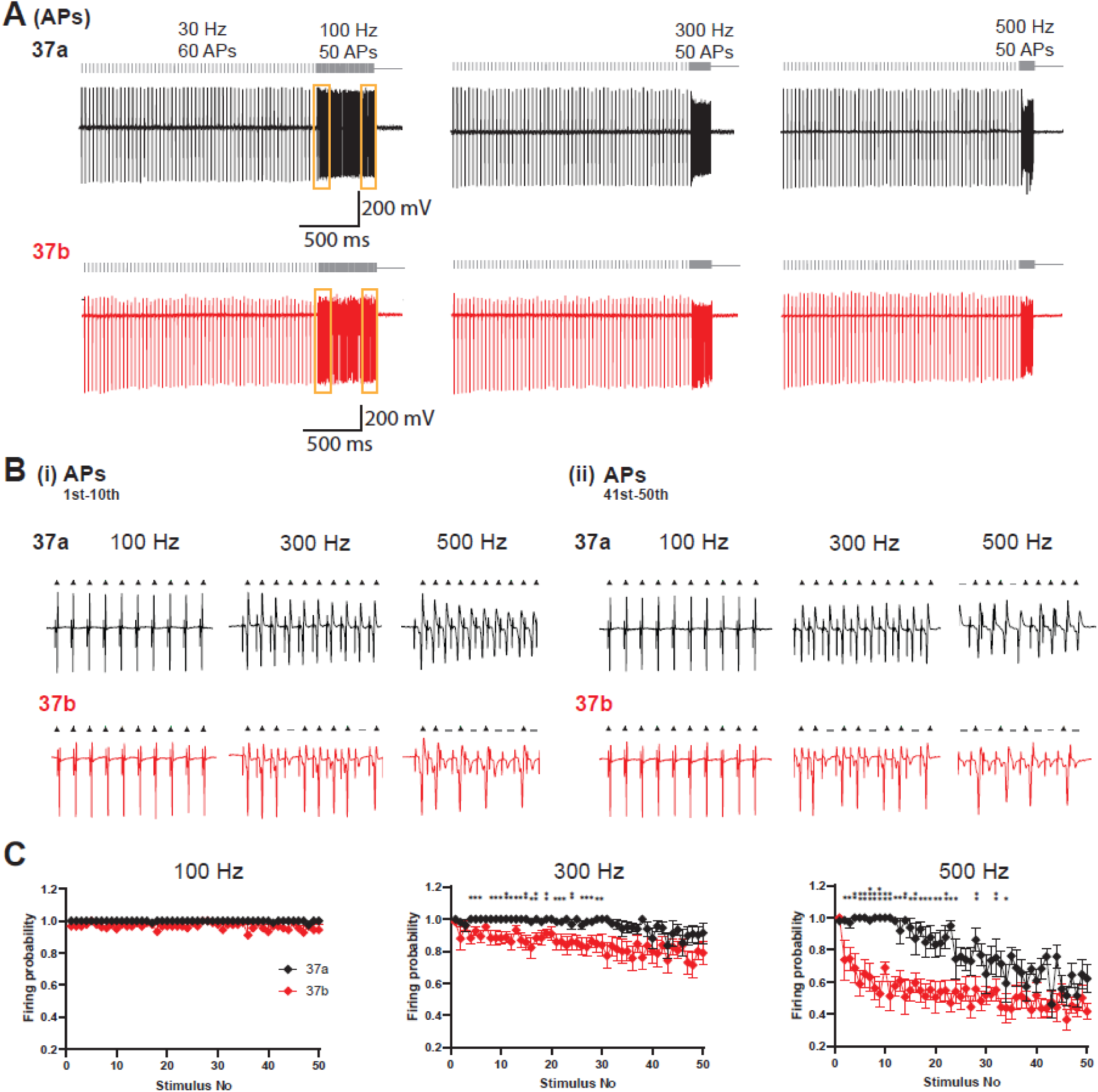
Lack of calcium current facilitation resulted in less firing probability of MNTB neurons. A: Representative traces of extracellular (cell-attached) spikes of MNTB neurons receiving calyces expressing Ca_V_2.1 37a (black, top) or Ca_V_2.1 37b (red, bottom). The failure of a spike is indicated by a gray flat line above the stimuli while the successful spike is indicated with a black triangle upward. Conditioned trains (60 AP) at 30 Hz preceding the tested trains (50 AP) at 100, 300, and 500 Hz. **B:** Expanded views of AP₁–_10_ (B-I) and AP₄₁–₅₀ (B-II) taken from the traces shown in panel **A**. **C: T**he firing probability plotted against the number of stimuli at different stimulus frequencies. Firing probability was calculated as the average number of spikes divided by the number of trials, 1 indicated no spike failure in all trials and 0 indicated no spikes at all.

In addition to AP reliability, the latency and jitter of the AP are key factors for the accurate encoding of auditory information. Therefore, we analyzed the jitter of the AP (standard deviation of spike timing ≥10 repetitions of each trial, see Methods). and latency (time determined from the start of stimulus artifact to the peak of spike (see Methods) at the different stimulation rates (**Fig 6**). We found increased latency of AP spikes in MNTB neurons innervated by Ca_V_2.1 37b calyces in a frequency dependent manner (**Fig 6B**). Spike latency showed consistent increases in latency in the Ca_V_2.1 37b calyces at both 100 Hz and 300 Hz. However, at 500 Hz there were clear differences in latency only in the first 10 AP at 500 Hz. Analysis of jitter showed that at 100Hz there was a small increase that was not consistent across the stimulus train. However, we found dramatic differences in jitter only in the first 35 APs at 300Hz, and only 15 APs at 500. This indicates that Ca_V_2.1 37b lacks the high temporal precision early in the stimulus, since this difference disappears later in the stimulus due to the increase in Ca_V_2.1 37a jitter. In addition, we found increased jitter, latency of AP, and decrease in synaptic firing probability in the “rested” Ca_V_2.1 37b calyx/MNTB synapses compared to the Ca_V_2.1 37a calyx/MNTB synapses (**Fig S6**) Based on these data, we conclude that CDF is important to post synaptic AP spiking reliability and precision at AP high firing rates.

**Figure 6.**
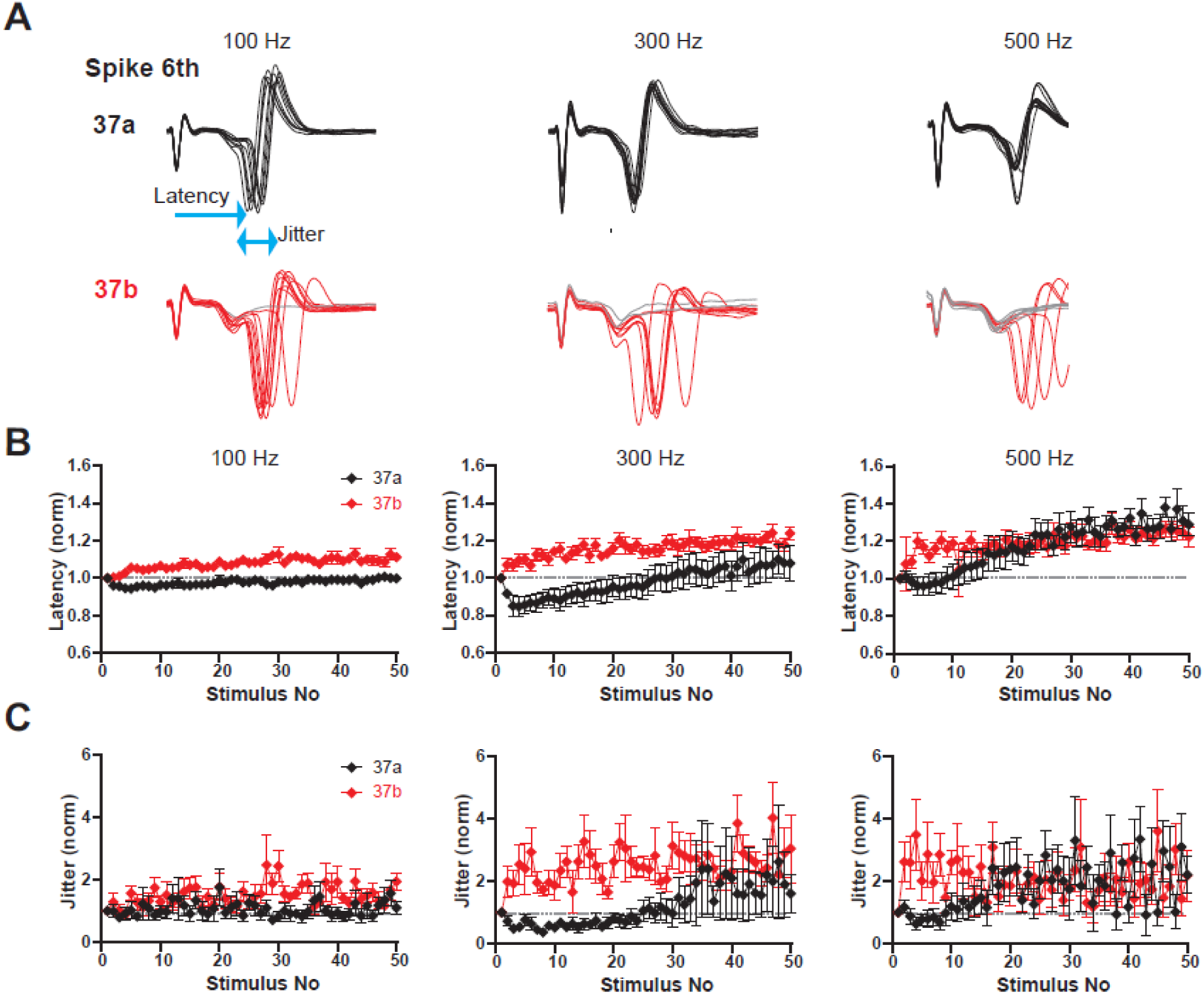
Loss of calcium current facilitation reduces the reliability of MNTB spikes and increases both latency and jitter. A: Overlay of a representative 6^th^ spike from 10 trials. Latency (single arrowhead) is defined as the time from the stimulus artifact to the peak of the spike. Jitter (double arrowheads) is defined as the standard deviation of spike timing across trials. The gray trace indicates a spike failure, while the preceding synaptic event remains present. **B:** Normalized spike latency, relative to the first spike, plotted as a function of stimulus number at different stimulation frequencies (100, 300, and 500 Hz) for Ca_V_2.1 37a (black) and Ca_V_2.1 37b (red) synapses. **C:** Normalized spike jitter, relative to the first spike, plotted as a function of stimulus number at different stimulation frequencies (100, 300, and 500 Hz) for Ca_V_2.1 37a (black) and Ca_V_2.1 37b (red) synapses. The gray dashed line at 1 indicates the baseline value (first spike) to facilitate comparison across stimulus trains. **Spikes Jitter** (2-10) at 100 Hz 37a; 0.80 ± 0.013, n=8, 37b; 0.13 ± 0.02 ms, n=9, p=0.04, at 300 Hz 37a; 0.80 ± 0.02, n=9, 37b; 0.19 ± 0.03 ms, n=10, p=0.005, at 500 Hz 37a; 0.06 ± 0.01 ms, n=6, 37b; 0.186 ± 0.01 ms, n=9, p=0.001. **Spikes Latency** (2-10) at 100 Hz 37a; 1.35 ± 0.04, n=8, 37b; 1.43±0.07 ms, n=9, p=0.4, at 300 Hz 37a; 1.3 ± 0.04 ms, n=10, 37b; 1.5±0.07 ms, n=10, p=0.04, at 500 Hz 37a; 1.33 ± 0.04, n=6, 37b; 1.52±0.04 ms, n=9, p=0.01.

### Mice with Ca_V_2.1 37b calyx of Held terminals have a reduced Auditory Brainstem Response Wave III amplitude

The MNTB provides well-timed inhibition to several nuclei in the superior olivary complex (SOC), including the LSO and MSO to encode ILDs and ITDs, respectively^19–22^. Since the calyx of Held is the sole excitatory input its ability to drive precise and reliable MNTB firing is essential for sound localization. Since the lack of CDF in the calyx of Held terminals resulted in decreased reliability and increased jitter and latency of MNTB AP spiking, we next asked how these deficits affected auditory processing *in vivo*.

To address this, we performed auditory brainstem response (ABR) recordings that measure the activity of neurons in response to sound stimuli from the cochlea to the CNS with Wave I representing AP from auditory nerve fibers leaving the cochlea and Wave III representing the strength of AP firing in the SOC. Since we expressed the Ca_V_2.1 splice variants in the calyx of Held that project to contralateral MNTB neurons, we expected that any changes in Wave III would reflect the ability of the calyx of Held to drive synchronous AP spiking in MNTB neurons. Therefore, we performed ABR recordings at increasing sounds pressure levels and frequencies with animals injected with either Ca_V_2.1 37a or Ca_V_2.1 37b and analyzed the ABR waves from animals P28 onwards (See Methods). Animals with little to no CN transduction were excluded from the analysis.

To first rule out if expression of the Ca_V_2.1 splice isoforms impacted synchronous AP firing in the CN, we first analyzed the Wave I amplitudes in the Ca_V_2.1 37a and Ca_V_2.1 37b mice. We found no differences in the Wave I amplitudes between Ca_V_2.1 37a or Ca_V_2.1 37b mice (**Fig 7A, B**). However, mice with Ca_V_2.1 37b calyces had smaller Wave III amplitudes, from sound pressure levels (SPL) from 55-100 dB SPL at 8 kHz and 16 kHz but not 32 kHz. (**Fig. 7A** red traces with arrowhead) when compared with those animals expressing Ca_V_2.1 37a (**Fig. 7A** black traces). 8kHz (at 80dB Ca_V_2.1 37a 1.9±0.22 μV, n=6, Ca_V_2.1 37b 0.685±0.08 μV, n=9, p= 0.0004) and 16kHz (at 80 dB Ca_V_2.1 37a 3.83±0.2 μV, n=6, Ca_V_2.1 37b 2.57±0.2 μV, n=9, p=0.001) but not 32kHz when compared with those mice expressing Ca_V_2.1 37a isoform in their calyces (at 80 dB Ca_V_2.1 37a 1.32±0.3, n=6, Ca_V_2.1 37b 1.7±0.2 μV, n=9,p=0.27) (**Fig. 7C**). Taken together, since this reduction was specific to Wave III with no change in Wave I, our data demonstrate that Ca_V_2.1 CDF is required for synchronous AP firing in the MNTB in response for sound localization. Therefore, we conclude that Ca_V_2.1 CDF is essential for reliable sound-evoked auditory brainstem transmission.

**Figure 7.**
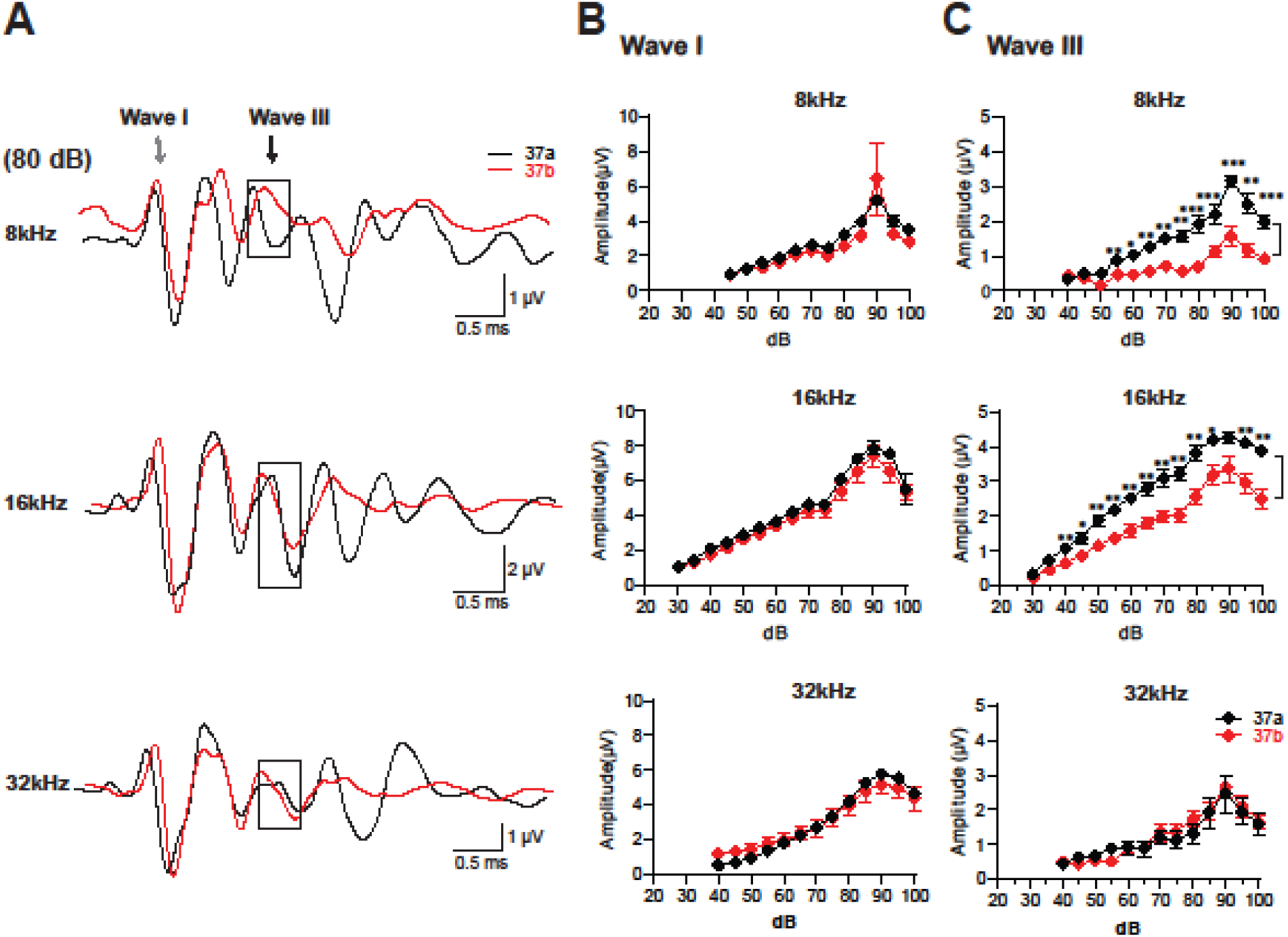
ABR analysis reveals reduced Wave III amplitude in Ca_V_2.1 37b mice compared to Ca_V_2.1 37a mice. A: Representative auditory brainstem response (ABR) waveforms recorded from live mice (postnatal day 28 or older) expressing either Ca_V_2.1 37a (black traces) or Ca_V_2.1 37b (red traces) splice variants. ABRs were measured across sound frequencies of 8, 16, and 32 kHz at intensities ranging from 25 to 100 dB SPL. Representative traces are shown at high sound intensity. Wave I (gray arrowhead) reflects activity of the auditory (cochlear) nerve and spiral ganglion neurons. Wave III (black arrowhead) corresponds to activity within the superior olivary complex (SOC), including the medial nucleus of the trapezoid body (MNTB). **B:** Wave I amplitude plotted as a function of sound intensity (dB SPL) for each frequency (8, 16, and 32 kHz; arranged from top to bottom). Wave I amplitude reflects the number of auditory nerve fibers activated, with larger amplitudes indicating recruitment of a greater neuronal population. The curves from both groups largely overlap, indicating that peripheral auditory nerve function was not affected by expression of either splice variant. **C:** Wave III amplitude plotted against sound intensity for each frequency. At 8 and 16 kHz, mice expressing Ca_V_2.1 37b exhibited consistently smaller Wave III amplitudes across a broad range of sound intensities compared to Ca_V_2.1 37a mice, with the difference becoming more pronounced at higher intensities. In contrast, no significant differences between splice variants were observed at 32 kHz. Wave III amplitude **8 kHz:** at 80dB 37a; 1.9±0.22, n=6, 37b 0.685 ± 0.08 μV, n=9, p= 0.0004), **16 kHz:** at 80 dB 37a 3.83 ±0 .2, n=6, 37b 2.57 ± 0.2 μV, n=9, p=0.001), **32 kHz:** at 80 dB 37a 1.32 ± 0.3, n=6, 37b 1.7 ± 0.2 μV, n=9,p= 0.27)

## Discussion

Elucidating the molecular mechanisms enabling high-fidelity computations in the auditory brainstem is essential for understanding the ability to accurate sound processing. By selectively expressing either the Ca_V_2.1 37a (CDF capable) or Ca_V_2.1 37b (CDF incapable) splice variant at the calyx of Held, we discovered that at physiological conditions Ca_V_2.1 CDF 1) results in activity dependent increases in presynaptic Ca^2+^ which is integral to synaptic facilitation; 2) is a key regulator of the reliability and temporal precision of post-synaptic AP firing, and 3) is crucial for synchronous AP firing in the SOC in response to sound. Therefore, we propose that presynaptic Ca_V_2.1 CDF is required for accurate auditory information processing.

### The role of Exon37 in Ca_V_2.1 abundance and AZ organization

Due to the steep Ca^2+^ dependence of AP mediated release, presynaptic Ca_V_2 abundance and coupling are key determinants regulating synaptic transmission and neuronal circuit output^45–47^. Although Ca_V_2.1 C-terminal AZ protein interaction sites are dispensable for regulating presynaptic Ca_V_2.1 abundance and coupling^36,48–50^, the Exon37 EF hand was proposed to be a key determinant setting presynaptic Ca_V_2.1 abundance and coupling^31^. Our data revealed no difference between Ca^2+^ current levels and initial EPSC amplitudes between the Ca_V_2.1 37a or Ca_V_2.1 37b calyx/MNTB synapses. Therefore, our data do not support a role for Exon 37 a/b splicing in controlling presynaptic Ca_V_2.1 abundance or AZ localization. Our data are similar to studies in primary cultured hippocampal neurons which demonstrated no difference in presynaptic Ca_V_2.1 37a or Ca_V_2.1 37b levels. However, these studies revealed a slight reduction in bassoon colocalization with Ca_V_2.1 37b, although EPSC amplitudes were similar between the Ca_V_2.1 37a and Ca_V_2.1 37b hippocampal neurons which suggests no difference in AZ localization.

Differences in coupling between SVs and Ca_V_2 channels underpin microdomain or nanodomain release^40,51^. The mature calyx of Held is a Ca_V_2.1 exclusive presynaptic terminal that utilizes nanodomain release (coupling ∼25 nm)^15,40–42^. Since we found no difference in the initial ESPC amplitude or waveform kinetics, our data do not support a role for the Exon37 EF hand in regulating coupling. This contrasts with previous conclusions that Exon37 splicing regulates coupling in hippocampal neurons as Ca_V_2.1 exon 37b dependent AP evoked release had more EGTA sensitivity but with no differences in initial AP evoked release between Ca_V_2.1 37a and Ca_V_2.1 37b^31^. However, these studies used the *CACNA1A* ΔExon47 cDNA while we used the full transcript (FT) *CACNA1A* cDNA. Prior work has shown that Ca_V_2.1 ΔExon47 has more mobility compared to Ca_V_2.1 FT that results in larger ESPC amplitudes but with higher EGTA sensitivity^48^. Thus, it was concluded that Ca_V_2.1 ΔExon47 synapses have higher release probability but looser coupling^48^. Therefore, it is possible that in the context of the Ca_V_2.1 ΔExon47 but not FT Ca_V_2.1, the different Exon37 splice variants adopt different Ca_V_2.1 conformations that impact or Ca_V_2.1 mobility and coupling. However, there are no known direct protein-protein interaction sites within Exon37 and there are no differences in the EPSC waveform kinetics with the Ca_V_2.1 IM-AA mutant^28–30^. Therefore, future experiments will be needed to further probe the mechanisms.

It is important to note that our phenotypes are not due to our HdAd rescue approach on the Ca_V_2.1 null calyx background. The use of the 470bp human synapsin promoter in the context of our transgene cassette in HdAd leads to similar expression level of calcium channels as wildtype calyces^36,50^ and we observed similar AP-evoked Ca^2+^ current amplitudes between our Ca_V_2.1 37a and Ca_V_2.1 37b calyces. Finally, we can rule out compensation by Ca_V_2.2 channels as no difference in synaptic transmission and plasticity was observed on either the *CACNA1A*^-/-^or *CACNA1A*^-/-^/*CACNA1B*^-/-^ background.

### Ca_V_2.1 CDF leads to activity dependent increases in presynaptic calcium and synaptic facilitation

Previous results at the immature calyx (P9-11) using whole cell paired recordings in 2mM external Ca^2+^ concluded that 40% of synaptic facilitation was due to CDF while the remaining 60% synaptic facilitation was due to build up residual free calcium^52^. We found Ca_V_2.1 37b calyces had negligible CDF and synaptic facilitation up to 500Hz. Thus, our findings at the mature calyx of Held show that at physiological temperature and external [Ca^2+^] that Ca_V_2.1 CDF is essential for synaptic facilitation. Differences between our findings and previous studies likely reflect the differences in the experimental approaches; square wave pulse 2 mM external calcium, developmental state, and room vs physiological temperature. Regardless, the mechanism by which Ca_V_2.1 CDF leads to synaptic facilitation such as to mobile or immobile buffer saturation^53^ or a separate facilitation sensor (Syt3 or Syt7)^54–56^ in the mature calyx remains to be studied. Finally, since the mature wild-type calyces and immature calyces exhibited activity dependent increases in presynaptic [Ca^2+^], we conclude that the Ca_V_2.1 splice variant found at the calyx of Held contains Exon37a at both immature and mature calyces of Held.

The role of Ca_V_2.1 CDF at other synapses is more controversial^28–31^. However, with respect to the role of Ca_V_2.1 CDF our data is similar to Ca_V_2.1 IM-AA mouse where significant reductions in synaptic facilitation were found at CA3/CA1 synapses and at the mouse NMJ with a reduced grip strength^29,30^. Our data and these studies differ with studies on Ca_V_2.1 IM-AA mouse at CA3-CA1, PF-PC, and PC/DCN synapses which concluded that Ca_V_2.1 CDF makes little contribution to synaptic facilitation at physiological temperatures and [Ca^2+^]^28^. However, there was a 50% reduction in Purkinje cell somatic Ca_V_2.1 currents. Since, presynaptic Ca^2+^ currents were not measured and these are global knock-in animals it is possible there were similar reductions in presynaptic Ca_V_2.1 currents which would impact CDF^32^ or a compensatory mechanism which results in reduced release probability at the synapse^28^. Furthermore, the differences may be due to differences in Ca_V_2.1 splice variants between synapses. Finally, our findings contrast with results at primary hippocampal neurons which used GCamp imaging to demonstrate that Ca_V_2.1 37b channels led to slight activity dependent increases in presynaptic Ca^2+^ and increased synaptic facilitation^31^. Given the diversity of synapses and the differences in the experimental conditions of these previous studies, future studies will be needed to unpack these differences.

### Ca_V_2.1 CDF and auditory signaling

The calyx of Held/MNTB synapse is a primary-like relay synapse that transforms well-timed excitation into well-time inhibition to multiple nuclei in the auditory brainstem and thereby controls monoaural and binaural sound processing^1^. Our results demonstrated that at 30 Hz and 100 Hz stimulation rates there was no difference in synaptic transmission between 37a or 37b terminals as there was no difference in ESPC amplitudes or postsynaptic firing. However, we found that at 300 Hz and 500 Hz stimulation rates the 37b synapses showed increased synaptic depression with a reduction in the steady state and decrease in reliability and temporal precision of post synaptic firing. Since reliability, latency, and jitter were significantly affected even at the 2^nd^ pulse, this indicates that Ca_V_2.1 CDF in the train stimuli acts as a band pass filter in activity dependent manner.

Since the RRP is not constant and changes in an activity dependent manner^57^, we hypothesize that Ca_V_2.1 CDF regulates the RRP to act as the bandpass filter. Previous studies demonstrated that ∼20-30 quanta are needed to trigger a postsynaptic AP^10^. Since the calyx of Held/MNTB synapse is tonically depressed, modeling simulations support a simple model of depression and recovery to maintain firing during activity dependent changes. At the calyx of Held and other fast firing synapses that activity dependent increases lead to faster SV recovery which are calcium dependent^9,58,59^. While we did not find an increase in the SV recovery rate in the conditioned synapses, we found that the cumulative EPSC (RRP) was smaller in 37b synapses compared to 37a synapses. Since SVs that are ∼25nm from Ca_V_2.1 are available for AP evoked release^41,43^, it is possible that activity dependent AP evoked increases in presynaptic Ca^2+^ entry manner expand the distance of SVs that remain accessible to AP evoked Ca²⁺ domains, potentially altering their release probability through SV priming, positional priming, or facilitation sensors. Regardless, it is important to note that our postsynaptic AP measurements did not contain kynurenic acid or NMDA blockers, therefore a direct correlation of the RRP size or RRP release kinetics between our voltage clamp experiments must be interpreted with caution as other mechanisms such as AMPAR receptor saturation may play a role.

Regardless of the molecular mechanisms, our findings demonstrate that loss of Ca_V_2.1 CDF resulted in a reduction in the ABR Wave III amplitude at 8kHz and 16 kHz. However, we did not see a difference at 32 kHz. Since we utilized C57BL/6 animals, which lose high frequency hearing it is possible that early cochlear degeneration at high frequencies masked the contribution of Ca_V_2.1 CDF at 32 kHz^60^. In this study, we used ABR to test monaural responses in the MNTB to sound stimuli, but did not assess binaural sound processing or other auditory processing pathways^61–65^. Since the MNTB innervates multiple nuclei, future experiments are needed to determine how the downstream pathways are impacted by the loss of Ca_V_2.1 CDF in the calyx. Regardless, our findings are consistent with prior studies demonstrating that dysfunction in calyx of Held/MNTB synaptic transmission leads to defects in auditory processing^62,66^.

Taken together, by selectively expressing either the Ca_V_2.1 37a or 37b splice variant at the calyx of Held, our findings provide new insights into how presynaptic Ca_V_2.1 CDF regulates high-fidelity computations for accurate auditory information processing.

## Acknowledgements

This work was supported by the National Institutes of Deafness and Other Communication Disorders (R01 DC014093 SMY, Jr., R01 DC018488 MH, and R01 DC012578 MH), the National Institute of Neurological Disorders and Stroke (R01 NS110742 SMY, Jr), the National Center for Advancing Translational Sciences (UM1TR004403 MH), and start-up funds from the University of North Carolina-Chapel Hill (S.M.Y., Jr). Deutsche Forschungsgemeinschaft (DFG, 420075000, CK)

We thank members of the Young lab for their comments on the manuscript. We thank Hao Li and Dr. Joe Zhou for help with HdAd production and titering. We thank Dr. Phillip Ng for HdAd packaging plasmids.

## Author contributions

M.A., J.L., C.K., B.H., carried out experiments and analyzed data. S.M.Y, Jr planned the project and analyzed data. M.H. analyzed data. M.A. and S.M.Y, Jr. wrote the manuscript. All authors jointly revised the paper. All authors approved the final version of the manuscript, agree to be accountable for all aspects of the work in ensuring that questions related to the accuracy or integrity of any part of the work are appropriately investigated and resolved, all persons designated as authors qualify for authorship, and all those who qualify for authorship are listed.

## Supplementary Information

Document: Supplemental Figures. Figure S1-S6

## STAR Methods

### Resource availability

#### Lead contact

Further information and requests for resources and reagents should be directed to the lead contact, Samuel M. Young, Jr.

### Materials availability

Transgenic mice used in this study are available from the lead contact upon request. This study generated new unique reagents which are the HdAd viral vectors.

### Data and code availability

All data reported in this paper will be shared by the <u>lead contact</u> upon request. Electrophysiological analysis software is available online: version 8.0.4.2, Wavemetrics, RRID:SCR_000325), Patcher’s Power Tools (version 1.19, RRID:SCR_001950), Neuromatic (RRID:SCR_004186). All data reported in this paper will be shared by the lead contact upon request.

## EXPERIMENTAL MODEL AND SUBJECT DETAILS

### Animal and ethical approval

All animals were housed either at the University of Iowa Animal Care Facility or the University of North Carolina-Chapel Hill animal facilities, approved under NIH assurance A3021-01 and certified by the State of Iowa and the State of North Carolina for use of living animals, and all experiments were done in accordance with the animal welfare laws of the University of Iowa and University of North Carolina-Chapel Hill Institutional Animal Care and Use Committee and Office of Animals Resources and complied with accepted ethical best practice. Mice were housed with a 12/12 hr light/dark cycle and ad libitum access to food and water. Post-natal day 1 (P1) mice were used for stereotaxic surgery, and experiments were done at P16 onwards except where noted.

## METHODS DETAILS

We used *Cacna1a fl/fl* mice or *Cacna1a fl/fl/ Cacna1b fl/fl* ^50^ of either sex at postnatal age P16-P23 for electrophysiology experiments and P28 onward for experiments performing the Auditory brainstem response (ABR) assay. P1 the mice were used for stereotaxic surgery.

### DNA Construct and Recombinant Viral Vector Production

Codon optimized (GENEart) Ca_V_2.1 α_1_ subunit cDNA Cacna1a (*Mus musculus*, Accession No. NP_031604.3), were used to generate Ca_V_2 α_1_ hybrid constructs. Cav2.1 37a and Cav2.1 37b constructs were sequenced verified. These constructs were individually cloned into the EcoRI and NotI sites of the 470 bp human synapsin (hSyn) neurospecific expression cassette ^67,68^.

### HdAd production

Helper-dependent Adenoviral vector (HdAd) (Cre) which co-expresses Cre recombinase (Cre and EGFP or mCherry under separate neurospecific expression cassettes driven by the 470 bp human synapsin promoter (hSyn) was produced as previously described ^67,68^. To produce HdAd Ca_V_2.1 37a and Cav2.1 37b viruses we followed standard protocols ^36,68^. Briefly, Ca_V_2 α_1_ cDNA constructs driven by hSyn promoter expression cassette were then cloned into he AscI site, pdelta23E4 which contains a separate neuro-specific expression of mCherry or myristolated EGFP (mEGFP) markers^69^. Production of HdAd was carried out as previously described ^68,70^. The HdAd plasmid was linearized with PmeI and then transfected (Profection Mammalian Transfection System, Promega, Madison, WI, USA) into 116 producer cells. Helper virus (HV) was added the following day. Forty-eight hours post infection, cells were subjected to three freeze/thaw cycles and lysate was collected. The HdAd vectors were amplified for total of five serial coinfections from 3×6 cm tissue culture dishes followed by one 15 cm dish and finally 30×15 cm dishes of 116 cells (confluence ∼90%). HdAd was purified by CsCl ultracentrifugation. HdAd was stored at −80ᵒC in storage buffer (10 mM Hepes, 1 mM MgCl_2_, 250 mM Sucrose, pH 7.4). HdAds were titered by as described previously ^50,71^

### Stereotaxic viral vector injections

We performed P1 stereotactic surgery as previously described^37,72,73^. Briefly, a Postnatal day 1 (P1) mice were anesthetized by a 5 min immersion into an ice bath. They were subsequently secured on a chilled aluminum block and checked throughout the surgery for anesthetic depth by tail pinch. Surgery was performed in an aseptic environment, using pulled pipettes with a 20 µm opening (Sutter Instruments, P1000), 2 µl of HdAd in storage buffer (containing in mM: 250 sucrose, 10 HEPES, 1 MgCl_2_ at a pH of 7.4) with 6.6% mannitol was injected into the Ventral Cochlear Nucleus (vCN) at a flow rate of 1 µl/min. For injection, we used a glass pipette ((Blaubrand, intraMark, catalog number: 708707)) filled with ∼ 2 μl of viral vectors (Helper-dependent adenovirus, HdAd) expressing mCherry-tagged iCre recombinase) along with EGFP-tagged Cav2.1 37/a or EGFP-tagged Cav2.1 37b (to express completely one splice variant in each pup), at a rate of 1 μl/min. The total amount of virus did not exceed 2*10^9^ viral particles. After 1-2 minutes of injection, the glass pipette was removed slowly to avoid the flow back of the injected solution. The pups then were taken to a recovery cage under a warm lamp 37 C° and returned to the mother’s cage after full recovery. If animals did not recover they were euthanized, and if they presented as distressed at any later timepoint veterinary staff were consulted, and their directions followed.

### Electrophysiology

#### Slice preparation

Acute brainstem slices were prepared as described previously^72^. briefly, after decapitation, brainstem was isolated and glued to a slicing chamber filled with ACSF containing the following (in mM): 125 NaCl, 2.5 KCl, 3 MgCl2, 0.1 CaCl2, 10 glucose, 25 NaHCO3, 1.25 NaH2PO4, 0.4 L-ascorbic acid, 3 myo-inositol, and 2 Na-pyruvate (pH 7.3–7.4, 310 mOsm). Coronal acute brainstem slices (150-250 μm in thickness) containing MNTB regions were obtained using a vibratome (VT1200, Leica Microsystems). Slices were transferred to an incubation beaker containing the extracellular solution (in mM): 125 NaCl, 2.5 KCl, 1 MgCl2, 1.2 CaCl2, 10 glucose, 25 NaHCO3, 1.25 NaH2PO4, 0.4 L-ascorbic acid, 3 myo-inositol, and 2 Na-pyruvate (pH 7.3–7.4, 310 mOsm) and was continuously bubbled with 95% O_2_–5% CO_2_ at 37 C°. Slices were incubated for ∼ 40-50 minutes for recovery before use.

In all experiments, slices were continuously perfused with aCSF and visualized by upright microscope (BX51W, Olympus) through 4X air or 60X water-immersion objective (LUMPlanFL N, Olympus, Tokyo, Japan) and a CCD camera (QI-Click, QImaging, Surrey, BC, Canada). Patch clamp recordings were performed by using an EPC 10/2 patch-clamp amplifier (HEKA, Lambrecht, Germany), operated by PatchMaster version 2×90.5 (Harvard Instruments, Holliston, MA, USA). Data were low-pass filtered at 6 kHz and sampled with a rate of 50 kHz. Calyces transduced with HdAd Ca_V_2.1 37a or 37b wre identified visually with two co-expressed mEGFP and mCherry markers. To visualize mEGFP and mCherry with light of 470 nm or 560 nm, respectively, a Polychrome V xenon bulb monochromator or a lumen Metal arc lamp was used (TILL Photonics, Grafelfing, Germany).

#### Afferent fiber stimulation-whole cell

Acute brainstem slices were transferred to the recording chamber containing the extracellular solution same as above. Slices were perfused with an oxygenated extracellular solution at a rate of 2-3 ml/min and physiological temperature ∼ 37°C. Glass pipettes were pulled using a horizontal Micropipette puller (Model P-1000, Sutter instrument company, USA) to get a pipette tip size (1-2 μm) and access resistance of (2-4 MΩ). For EPSC recording, pipettes were filled with internal solution containing the following (in mM): 130 Cs-gluconate, 20 tetraethylammonium (TEA)-Cl, 10 HEPES, 5 Na2-phosphocreatine, 4 Mg-ATP, 0.3 Na-GTP, 6 QX-314, and 5 EGTA (pH 7.2, 318 mOsm). Pipette resistance was compensated online to 3 MΩ and offline to 0 MΩ.

Afferent fibers of the calyx of Held terminals were evoked electrically using parallel bipolar platinum-Iridum microelectrodes (# 221922, FHC Inc., USA) placed halfway between the midline of the brainstem and MNTB. Simulation voltage of 0.1-5V was produced by stimulus isolator (ISO-Flex, GmbH, Germany) to stimulate the afferent fibers. EPSCs were recorded from MNTB neurons in the presence of 0.25 mM kynurenic acid to prevent postsynaptic AMPA receptors saturation, 50 μM of D-AP5 to block NMDA receptors, 20 µM bicuculline to block GABA_A_ receptors, and 5 µM strychnine to block glycine receptors (all Tocris Bioscience). Postsynaptic MNTB principal neurons were whole-cell voltage-clamped at −60 mV, and R_S_ (<8 MΩ) was online compensated to <3 MΩ (EPC10/2 amplifier, PatchMaster 2.90.5, HEKA Elektronik). We recorded neurons with or without a conditioning protocol for conditioned stimulation, we used conditioned train at 30 Hz for ∼ 2S followed by test train at 100 Hz,300 Hz, and 500 Hz) with at least 3 repetitions for each trial and recovery period (30-60S). The conditioning frequency of 30 Hz was chosen to mimic the in vivo condition of spontaneous activity of MNTB neurons (^74^).

#### Loose patch (Cell-attached) recordings

To record extracellular spikes of MNTBs, recording pipette same as above (tip size 1-2 μm) was filled with ACSF to avoid impacting the receptors and ion channels located on the cell membranes of recorded cells and in the presence of inhibitory blockers (5 μM strychnine) and (20 μM bicuculline). Extracellular spikes were recorded in voltage clamp configuration with holding potential 0 mV and physiological temperature ∼ 37°, and after series resistance reached 40-100 MΩ. The afferent fibers were induced using parallel bipolar electrodes (conditioned train, 30 Hz for ∼ 2S followed by test train, 100 Hz, 300 Hz, and 500 Hz) with 10-13 repetitions for each trial. Cells with the break in membranes were excluded from analysis, this can be detected when one phase of spike responses are gone (the outward phase). We defined spike failure as occurring when a neuron received a weak input (subthreshold response) that was not strong enough to trigger a spike. This subthreshold response was typically observed as a small synaptic event that preceded spikes. Stimulation failure, in contrast, occurred when electrical stimulation failed to evoke any synaptic response. neurons showing stimulation failure in any pulse of a stimulus train were excluded from the analysis.

#### Whole-cell presynaptic Ca^2+^ current recordings

To isolate presynaptic Ca^2+^ currents, aCSF was supplemented with 1 or 1.5 μM tetrodotoxin (TTX, Alomone labs), 100 μM 4-aminopyridin (4-AP, Tocris) and 20 mM tetraethylammonium chloride (TEA, Sigma Aldrich) to block Na^+^ and K^+^ channels. Calyces were whole-cell voltage clamped at −80 mV, all presynaptic recordings were acquired at physiological Ca^2+^ (1.2 mM) and temperature (37°C) to mimic *in vivo* condition. 100 μM 4-AP (Tocris Bioscience), 20 mM TEA (Sigma Aldrich) and 1 μM TTX (Alomone Labs) were used to isolate Ca^2+^ currents. Presynaptic patch pipette with open tip diameters 4-6 MΩ resistance were pulled from 1.5 mm thin-walled borosilicate glass (Harvard apparatus) and were filled with same solution as the postsynaptic recording pipettes with the exception with that 0.1 mM EGTA was used and QX-314 was excluded. To determine total presynaptic Ca^2+^ current was voltage-clamped at −80 mV and depolarized to 10 mv with square pulse for 10 ms duration. To test for CDF, the presynaptic terminal was clamped at ramp pulse to +60 mV for 1 ms duration with rising time of 0.2 ms and falling time of 0.5 ms^44,75^ at different simulation frequencies (100, 300, and 500 Hz). Pipettes were coated with dental wax to minimize stray capacitance and improve voltage clamp quality. Presynaptic series resistance was less than 15 MΩ and was compensated online to 3 MΩ or less with a time lag of 10 µs by the HEKA amplifier. Leak and capacitive currents were subtracted online with a P/5 routine. Cells with series resistance >15 MΩ and leak currents > 150 pA were excluded from the analysis.

#### Auditory brainstem response (ABR) assay

Mice of either sex (P28∼) are anesthetized with an injectable ketamine + xylazine cocktail (87.5 mg/kg ketamine/ 12.5mg/kg Xylazine) and placed in an audiometric booth. Acoustic stimuli are presented to the ear (RZ6 multi-IO/processor, BioSig RZ version 5.7.6 software, Tucker-Davis Technologies (TDT)) via MF1- M mono speaker (TDT) coupled to the ear via a speculum inserted into the external auditory canal. ABRs are measured utilizing the Medusa 4Z bioamp (TDT) and three subdermal needle electrodes (Lifesync, product# NS-S83018-R9-10) that are placed in the head (vertex), ipsilateral mastoid, and mice back (ground electrode). The responses were then averaged over 512 presentations for each auditory stimulus (100-20 db SPL) for 8KHz,16KHz, and 32KHz.

## QUANTIFICATION AND STATISTICAL ANALYSIS

### Electrophysiological analysis

All electrophysiological data was analyzed using Igor Pro (version 8.0.4.2, Wavemetrics, RRID:SCR_000325) equipped with Patcher’s Power Tools (version 2.19, RRID:SCR_001950) and NeuroMatic (RRID:SCR_004186)^76^.

electrophysiological and ABR data were analyzed offline using a custom built-in function (Neuromatic v3.01)(^76^) equipped in Igor Pro (version 8.0.4.2, Wavemetrics, RRID:SCR_000325).

EPSC amplitude was determined as the peak of EPSC minus baseline preceding stimulus artifact. Rise time was measured as the time between 10%-90% of EPSC amplitude, full width half maximum (FWHM) as the width of EPSC measured at 50% of its amplitude, and onset delay as the time between stimulus artifact and the maximum curvature point of EPSC rising flank fitted with a Boltzmann equation (^77^). Jitter of extracellular spikes (standard deviation of spikes timing) was determined using 10-13 repetitions of each trial and with at least 6 successful responses from those showing high failure rates (usually at 500 Hz), latency of spikes as the time between stimulus artifact and the negative peak of extracellular spikes and firing probability as the number of spikes divided by the stimulus numbers.

Calcium currents amplitude was measured from baseline to peak. For AP waveform presynaptic Ca^2+^ currents, each amplitude of AP train was normalized to the 1^st^ spike amplitude and fitted against the spike number.

### ABR data analysis

ABR data were analyzed offline using a custom built-in function (Neuromatic v3.01)(^76^) equipped in Igor Pro (version 8.0.4.2, Wavemetrics, RRID:<u>SCR_000325</u>). ABR wave III amplitude was measured from the trough to the peak of Wave III.

### Statistics

Data were presented as a mean ± SEM (*n* = number of cells). Normality of data and equality of variance were evaluated by Kolmogorov-Smirnov test and *F* test, respectively. Statistical significance was determined by unpaired t-test (two-tailed) for comparison between two groups if they passed normality and equality of variance tests or Mann-Whitney test if they did not pass the normality test. Unpaired t-test with Welch’s correction was used to compare two groups which have unequal variances. The level of significance was set at 0.05.

