## Supplemental figures for "Presynaptic Ca_V_2.1 calcium dependent facilitation is essential for faithful auditory information transfer"

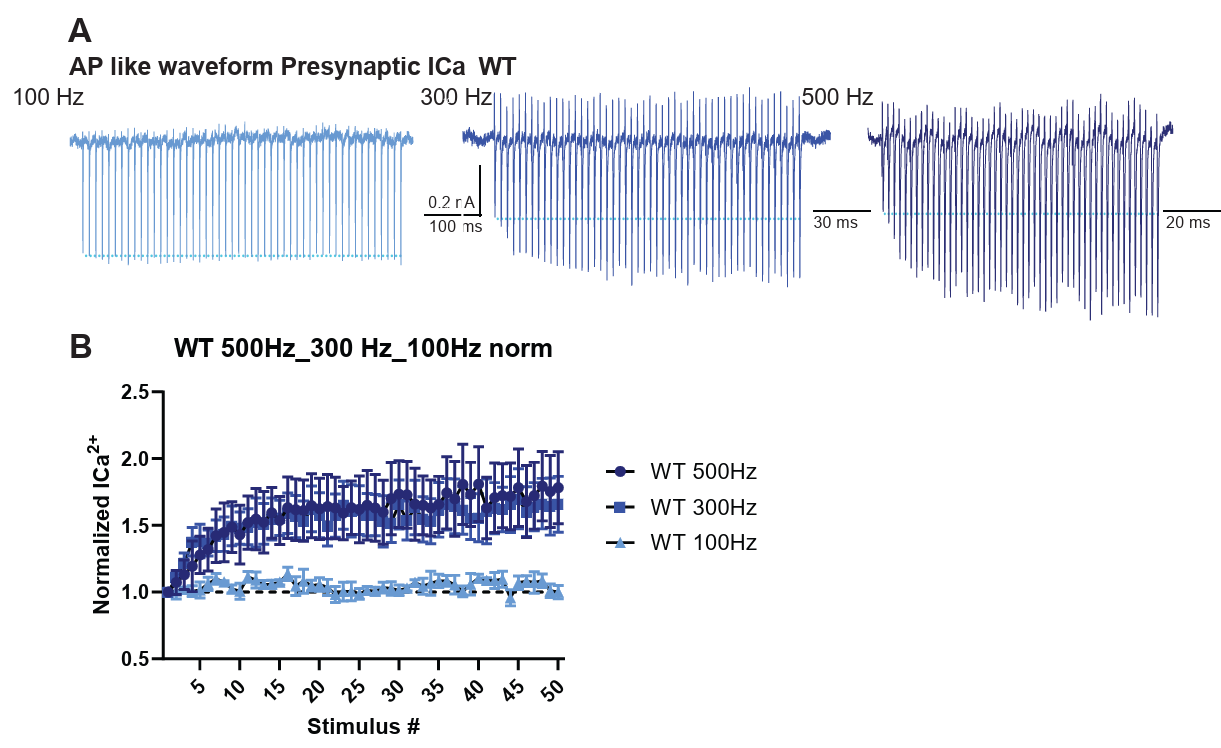

**Figure S1. Wild-type calyces exhibit robust Ca²⁺-dependent facilitation at high frequencies.**

**A:** Representative presynaptic whole-cell recordings of AP like waveform train (100, 300, 500 Hz) at p16-19 wildtype calyces. **B**: Graphs of Ca^2+^ dependent facilitation, Ca^2+^ current amplitudes are normalized to the 1st spike and plotted against stimulus number at different frequencies (100, 300, 500 Hz). n=4, 4 animals for each frequency. Related to Figure 2.

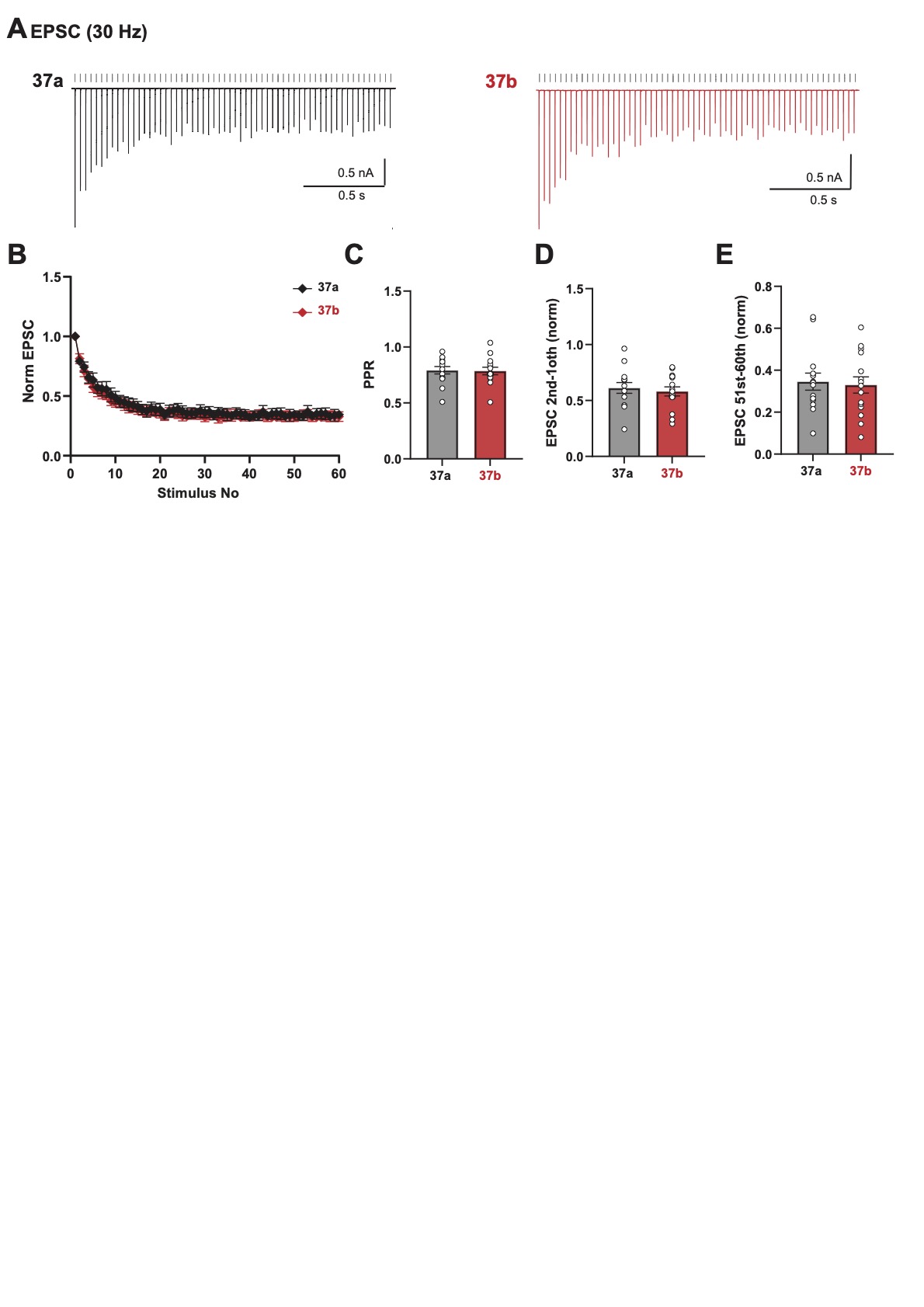

**Figure S2. Lack of Ca²⁺ current facilitation does not affect short-term synaptic plasticity during in vivo–like spontaneous activity.**

**A:** Representative postsynaptic whole-cell recordings from principal MNTB neurons receiving calyces expressing Ca_V_2.1 37a (black) or Ca_V_2.1 37b (red). EPSCs were evoked at 30 Hz, a frequency within the range of in vivo–like spontaneous firing. **B:** EPSC amplitudes normalized to the first response and plotted as a function of stimulus number for Ca_V_2.1 37a (black) and Ca_V_2.1 37b (red). Both groups exhibited a comparable pattern of short-term synaptic depression. **C:** Paired-pulse ratio (PPR) analysis revealed no significant difference between Ca_V_2.1 37a and Ca_V_2.1 37b synapses. **D:** Normalized steady-state EPSC amplitudes (EPSC 41–50 relative to the first EPSC) were not significantly different between neurons receiving Ca_V_2.1 37a- or Ca_V_2.1 37b-expressing calyces. Related to Figure 3.

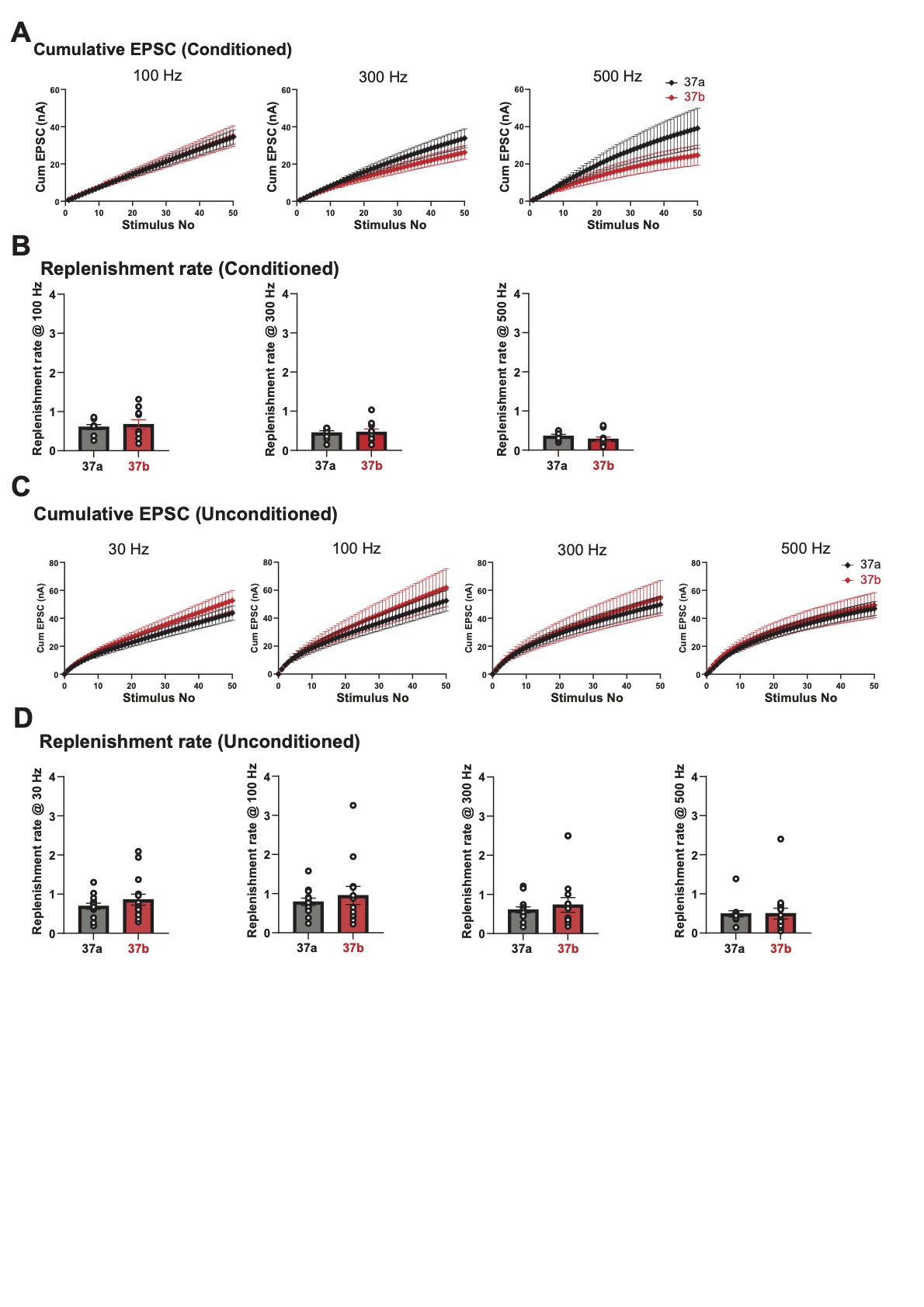

**Figure S3. Lack of Ca²⁺ current facilitation does not alter EPSC replenishment rates.**
**A:** Cumulative EPSC plots from test trains (100, 300, 500 Hz) following 30 Hz conditioning, plotted against stimulus number, showing a frequency-dependent increase in cumulative EPSCs. **B:** Comparison of replenishment rates between 37a and 37b synapses, showing no significant differences. **C:** Cumulative EPSCs from unconditioned trains plotted against stimulus number for both 37a and 37b synapses. **D:** Replenishment rates at unconditioned stimulus frequencies (30, 100, 300, and 500 Hz), showing no statistically significant differences between the two splice variants. Related to Figure 3 and 4.

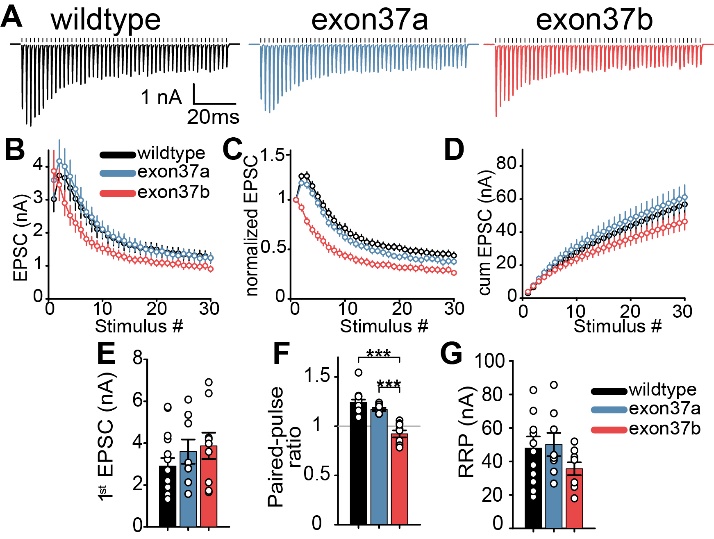

**Figure S4. Exon37a is required for synaptic facilitation A)** Representative traces AP-evoked EPSCs at 500 Hz on Cacna1a/Cacna1b null background. **B)** EPSC amplitudes **C)** Normalized EPSC amplitudes **D)** cumulative EPSC **E**) 1^st^ EPSC amplitude **F**) paired pulse ratio (P2/P1) **G**) RRP size (*wildtype* n=14, *exon37a* n= 8, *exon37b* n=9) ***p<0.001, One Way ANOVA

Related to Figure 4.

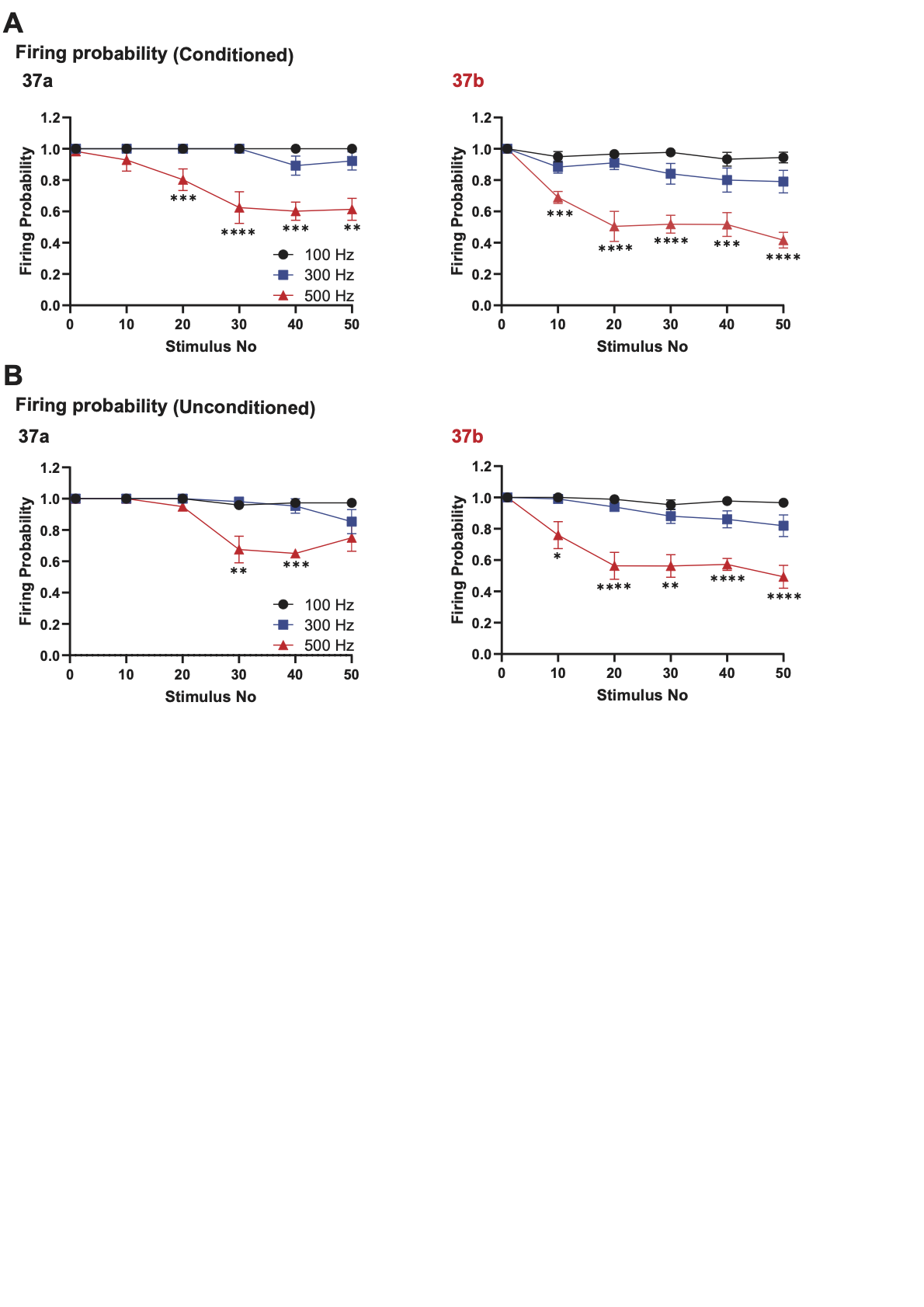

**Figure S5.**

**Distinct frequency-dependent firing probability patterns in MNTB neurons receiving 37a versus 37b calyceal inpu**ts**.**

**A:** Representative loose-patch recordings from principal MNTB neurons. The firing probability at different stimulation frequencies (100, 300, and 500 Hz), preceded by a conditioning stimulus of 30 Hz for 60 action potentials (APs), is plotted against stimulus number (50 stimuli). Frequency-dependent changes in firing probability were observed in both groups. In neurons receiving **37a** calyces, this effect became significant in the later part of the stimulus train (around stimulus 20 and thereafter) at 500 Hz compared with 100 and 300 Hz. In contrast, neurons receiving **37b** calyces exhibited significant frequency dependence earlier in the stimulus train (starting from approximately stimulus 10 and onwards). **B:** Unconditioned firing probability plotted against stimulus number (50 stimuli). MNTB neurons receiving **37a** calyces showed significant frequency dependence only during the later part of the stimulus train (stimuli ~20–30). In contrast, neurons receiving **37b** calyces displayed stronger frequency dependence beginning earlier in the stimulus train (approximately stimulus 10) and continuing until the end of the train (stimulus 50). Related to Figure 5.

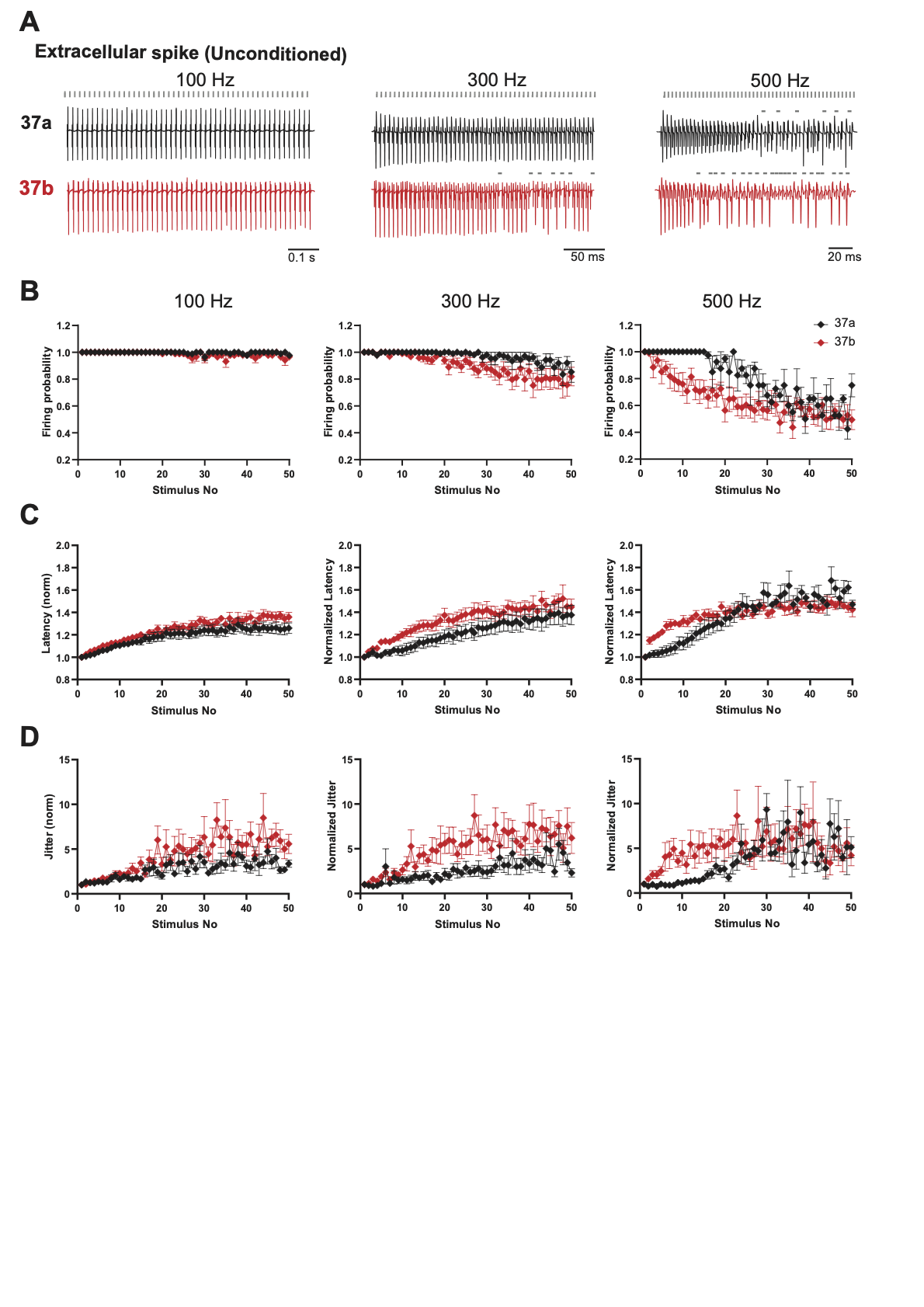

**Figure S6. Reduced spike reliability in MNTB neurons receiving Ca_v_2.1 37b calyceal inputs during high-frequency stimulation.**

**A:** Representative unconditioned extracellular spikes (50 APs) recorded in cell-attached mode from MNTB neurons receiving Cav2.1 37a synapses (black, top) or Cav2.1 37b synapses (red, bottom) at stimulus frequencies of 100, 300, and 500 Hz. Gray dashed lines above the traces indicate spike failures. **B:** Firing probability plotted as a function of stimulus number at each frequency. At 100 Hz, firing probability was comparable between Cav2.1 37a and 37b synapses. At 300 Hz, Cav2.1 37b synapses exhibited more spike failures during the steady-state phase compared to Cav2.1 37a synapses. At 500 Hz, occasional spike failures were observed in the steady state of Cav2.1 37a synapses; however, failures were more frequent and occurred earlier during stimulation at Cav2.1 37b synapses. **C–D:** Latency and jitter were increased at Cav2.1 37b synapses, particularly during the early phase of stimulation at 300 and 500 Hz. No significant differences were observed at 100 Hz. These findings suggest a critical role for CDF in maintaining spike fidelity during high-frequency transmission. Related to Figure 5, 6.
